# Using oxygen and hydrogen stable isotopes to track the migratory movement of Sharp-shinned Hawks (*Accipiter striatus*) along Western Flyways of North America

**DOI:** 10.1101/856278

**Authors:** Elizabeth A. Wommack, Lisa C. Marrack, Stefania Mambelli, Joshua M. Hull, Todd E. Dawson

**Author notes:** Tropical Conservation Biology and Environmental Science (**TCBES**) Graduate Program, University of Hawaii, Hilo, Hawaii, United States of America (Lisa C. Marrack). Corresponding author, (EAW). These authors contributed equally to this work (Elizabeth A. Wommack, Lisa C. Marrack, Stefania Mambelli).

## Abstract

The large-scale patterns of movement for the Sharp-shinned Hawk (*Accipiter striatus*), a small forest hawk found throughout western North America, are largely unknown. However, based on field observations we set out to test the hypothesis that juvenile migratory *A. striatus* caught along two distinct migration routes on opposite sides of the Sierra Nevada Mountains of North America (Pacific Coast and Intermountain Migratory Flyways) come from geographically different natal populations. We applied stable isotope analysis of hydrogen (H) and oxygen (O) of feathers, and large scale models of spatial isotopic variation (isoscapes) to formulate spatially explicit predictions of the origin of the migrant birds. Novel relationships were assessed between the measured hydrogen and oxygen isotope values of feathers from *A. striatus* museum specimens of known origin and the isoscape modeled hydrogen and oxygen isotope values of precipitation at those known locations. We used these relationships to predict the origin regions for birds migrating along the two flyways from the measured isotope values of migrant’s feathers and the associated hydrogen and oxygen isotopic composition of precipitation where these feathers were formed. The birds from the two migration routes had overlap in their natal/breeding origins and did not differentiate into fully separate migratory populations, with birds from the Pacific Coast Migratory Flyway showing broader natal geographic origins then those from the Intermountain Flyway. The methodology based on oxygen isotopes had, in general, less predictive power than the one based on hydrogen. There was broad agreement between the two isotope approaches in the geographic assignment of the origins of birds migrating along the Pacific Coast Flyway, but not for those migrating along the Intermountain Migratory Flyway. These results are discussed in terms of their implications for conservation efforts of *A. striatus* in western North America, and the use of combined hydrogen and oxygen stable isotope analysis to track the movement of birds of prey on continental scales.

## Introduction

Thousands of bird species migrate, traveling from breeding territories to wintering grounds and back annually, sometimes over vast distances and geographical features [1]. Such seasonal movements make the factors that stress populations of migratory birds difficult to track, as individuals may be affected by changes at any point along their migratory route [2–4]. As a result, understanding the migratory paths and connections between breeding and wintering sites is critical for strategizing conservation and preservation actions for particular bird species, such as raptors which are often secretive and breed over a wide area [5].

Migratory routes taken by different birds that follow similar pathways across continents are identified as migratory “flyways”, which are hypothesized to represent the shortest and least costly course over wide geographic distances [6]. Differences in the choice of migratory flyway between and within species may represent evolutionary divergences related to specific adaptations [7–9]. For some species, individuals that stray from or cross between flyways experience higher costs to migration, therefore evolutionary associations are likely to exist between populations of birds and specific migratory flyways [10–12]. One common way to monitor populations of birds of prey is by tracking migratory movement at watch sites and banding stations along migratory flyways [5, 13, 14]. Long-term data sets from raptor migration watch-sites can indicate population dynamics for specific species and populations, which can provide indications of changes in population size [15–19]. However, it can be difficult to connect migrating birds to other geographic stages in their life cycle (breeding and wintering) through these data, and without these connections it may be problematic to understand what factors underlie population size changes [20].

The Sharp-shinned Hawk (*Accipiter striatus*) is a small, forest hawk found throughout North America [21]. During breeding, *A. striatus* prefers dense coniferous and deciduous forests, making nesting sites difficult to identify and survey, and breeding birds problematic to monitor [21–23]. Heavy persecution resulting from the shooting of thousands along migration routes each year in the late 19^th^ and early 20^th^ centuries [22, 24], combined with the effects of dichlorodiphenyltrichloroethane (DDT) used as a pesticide in the early 20^th^ century, resulted in *A. striatus* numbers decreasing steadily across the species’ range [25]. However, over the past several decades, data from population and migratory monitoring of the species have shown inconsistent and contrary trends of both significant increases and decreases [26, 13]. Since North America’s smallest hawk is difficult to track on its breeding grounds, most population monitoring occurs at watch and banding sites along migratory flyways. Connecting migratory flyways and watch sites with specific breeding ranges will allow for a greater understanding of wider populations trends for this secretive bird of prey.

Raptors in the western side of the North American continent are believed to travel along three migratory flyways: the Pacific Coast, the Intermountain, and the Rocky Mountain Flyways [12]. Band recovery data of *A. striatus* trapped at migration sites along each of the flyways have shown demographic population differentiation along each route [12, 15, 27]. However, sample sizes for band returns are often small and may therefore represent biased information [28]. Endogenous markers, such as genetic analysis and stable isotope analysis (SIA), are not able to provide the same level of precision on locality as extrinsic markers (i.e. bands, satellite telemetry), but are easier and less expensive to use with large sample sizes, and can provide a larger return in data for wide geographic areas [29–35]. Integrating information from both levels of analysis can provide the best account of current population trends, and allow researchers to connect migratory flyway monitoring data with information on geographical breeding areas.

Over the past few decades, a number of studies have used SIA of naturally occurring hydrogen (H) of feathers in order to estimate migratory patterns and ecological connectivity among habitats for a broad range of species of birds [36]. In comparison, only recently with advances in continuous flow pyrolysis techniques have reliable SIA of oxygen (O) of organic materials been possible. Therefore, much less data exist on the additional information that can be obtained by measuring both elements to trace animal movements from different tissue types, such as feathers and hair [37–40].

Keratin, the main constituent of feathers, remains chemically inert following its synthesis, and it incorporates hydrogen and oxygen from consumed water and dietary sources. The stable hydrogen and oxygen isotopic compositions in the keratin of a feather (δ^2^H_F_ and δ^18^O_F_ values) therefore become markers of the environmental conditions and will vary geographically due to the spatial variation in the stable hydrogen and oxygen isotopic composition of meteoric precipitation (δ^2^H_P_ and δ^18^O_P_ values) [41]. This variation is caused primarily by isotope effects associated with evaporation and condensation processes [42–44] and correlates inversely with latitude and elevation across the continents [45]. Additionally, because hydrogen and oxygen are incorporated or “routed” into organic compounds like keratin along different metabolic pathways, the δ^2^H and δ^18^O of feather keratin may not provide the same information even if these elements had their origin from the same water source.

Linking the isotopic composition of animal tissue to geographic locations of origin is based on important assumptions, such as the presence of known and constant isotope effects associated with tissue synthesis, and an understanding of the relationship between the isotopic compositions of the tissue and that of the environmental food web and water source signals [46, 39, 74]. However, work on some species of birds suggests a decoupling between hydrogen and oxygen isotope compositions in food webs that might affect the usefulness of δ^18^O_F_ measurements for assignment of bird origin [48, 49, 39].

Previous SIA studies have used δ^2^H_F_ values to estimate the timing and pattern of migration for *A. striatus* from western North America [50, 51]. Birds migrating along the Intermountain Flyway were found to originate from the Northern Rocky Mountain Range (Idaho (ID) and Montana (MT)) in the United States of America (USA) north through British Columbia (BC, Canada) [51]. Specific origins were not determined for birds caught along the Rocky Mountain Flyway, but it has been shown that *A. striatus* followed a chain migration pattern where individuals from lower latitudes migrated earlier than those from higher latitudes [50]. However, no work has been done to look at the origins of *A. striatus* from the Pacific Coast Flyway or has incorporated two isotopes to trace the origin of birds from neighboring flyways.

Migration can be a difficult and dangerous behavior for birds, and specific adaptations for breeding populations that use different flyways would be anticipated to lead to population differentiation [52]. If flyways represent unique migratory paths for different populations, then it can be predicted that *A. striatus* migrating along neighboring flyways should originate from separate breeding origins, as hinted at by their band results [12, 15]. However, no genetic difference in mitochondrial control region sequences has been found between *A. striatus* caught on migration along either flyway [53], suggesting that either birds that utilize the different routes overlap and intermix in their breeding populations or that mitochondrial markers lack the variability to resolve population genetic differentiation among flyways. Use of nuclear microsatellite loci has demonstrated differentiation between the intermountain and Pacific flyways in another wide-ranging raptor, the Red-tailed Hawk (*Buteo jamaicensis*) [54], suggesting that similar differences among flyways may exist in *A. striatus*.

In this study, we used the variation in δ^2^H_F_ and δ^18^O_F_ values of *A. striatus* and large scale models of spatial isotopic variation (isoscapes)for hydrogen and oxygen to: a) examine the origin of *A. striatus* caught along the Pacific Coast Flyway in comparison to birds caught along the Intermountain Flyway, and b) investigate the usefulness of oxygen isotopes to determine the origin of raptorial birds of prey. First, we established relationships, separately for hydrogen and oxygen, between δ^2^H_F_ and δ^18^O_F_ values of *A. striatus* museum specimens from known natal locations, to predict the isotope values of precipitation at the origin regions of birds migrating along the Pacific Coast and Intermountain Flyways from the measured isotope values of migrant’s feathers. Prediction of precipitation hydrogen and oxygen isotope composition and assignment of migrating bird origin locations were accomplished using an internet-based environmental water isoscape of Western North America (http://isomap.org) [55]. Finally, predictions of sites of origin based on δ^2^H_P_ and δ^18^O_P_ values were also compared.

Assuming that the choice of migratory flyway is driven by evolutionary processes [11], and that it would be costly for individuals to jump between migratory routes, we predicted that *A. striatus* caught along the Pacific Coast Flyway would originate from further west then those caught along the Intermountain Flyway, and show little overlap in their determined natal origins. However, instead the SIA revealed that birds from the Pacific Coast Flyway origins extended further east then expected, including into the Rocky Mountain Range and the western interior of North America. We report that both hydrogen and oxygen isotopic analysis with feathers can be used to determine the origin of migratory birds of prey, but that caution must be taken when interpreting the outcome.

## Materials and methods

### Sample collection

Contour feathers of *A. striatus* from the ventral sternal feather track (*n* = 23 juveniles (Table 1, Fig 1) and *n* = 25 adults (Table S1, S1 Fig) were sampled from museum specimens of known collection locality. *A. striatus* follow a complex basic strategy of molt, and have a limited to absent preformative molt which occurs for hatch-year and second-year birds between December and April of their first year. Adult molt (the definitive prebasic molt) occurs primarily on breeding grounds, starting during egg laying and incubation [21, 56]. As a result of this molt strategy both juvenile and adult *A. striatus* contour feathers have a high probability of being grown either in the location where they were hatched or where they were breeding. Contour feathers in ventral sternal feather track were selected for the small size of the feather, the ease of repeatability of selection of feathers from the same track for collection from individual birds, and as contour feathers have been used successfully to track movement of *A. striatus* in previous work [51]. Feather samples of *A. striatus* were obtained from the following museums in the USA: California Academy of Sciences (CAS), San Francisco, CA; Charles R. Conner Museum, Pullman (CRCM), WA; Museum of Vertebrate Zoology (MVZ), University of California, Berkeley, CA; San Diego Natural History Museum (SDNHM), San Diego, CA; University of Wyoming Museum of Vertebrates (UWYMV), Laramie, WY. In order to acquire samples that were representative of feathers grown on breeding and natal geographic regions of interest, specimens were constrained by date of collection (15^th^ March – 30^th^ August), and feather wear [56, 21, 57]. Only feathers with little or no wear were from museum specimens were selected to guarantee that they were grown in the present year and at the location of collection. Feathers were also sampled from migrating juvenile *A. striatus* banded at the Marin Headlands, Marin County, CA (Golden Gate Raptor Observatory (GGRO), Pacific Coast Flyway, *n* = 20), and the Goshutes Mountains, Elko County, NV (HawkWatch International (Goshutes), Intermountain Flyway, *n* = 10) between August and October 1999 (see [53] for sampling protocol). The migratory birds used in this study were collected with samples previously used to analyze the genetic population structure of *A. striatus*, and so are known to be from the western genetic population [53]. However, the specific origin of migrating individuals was unknown. No live birds were handled for the described study. Sampling of birds at banding stations was performed in 1999, and feathers used in this study were from archived collections.

**Fig 1.**
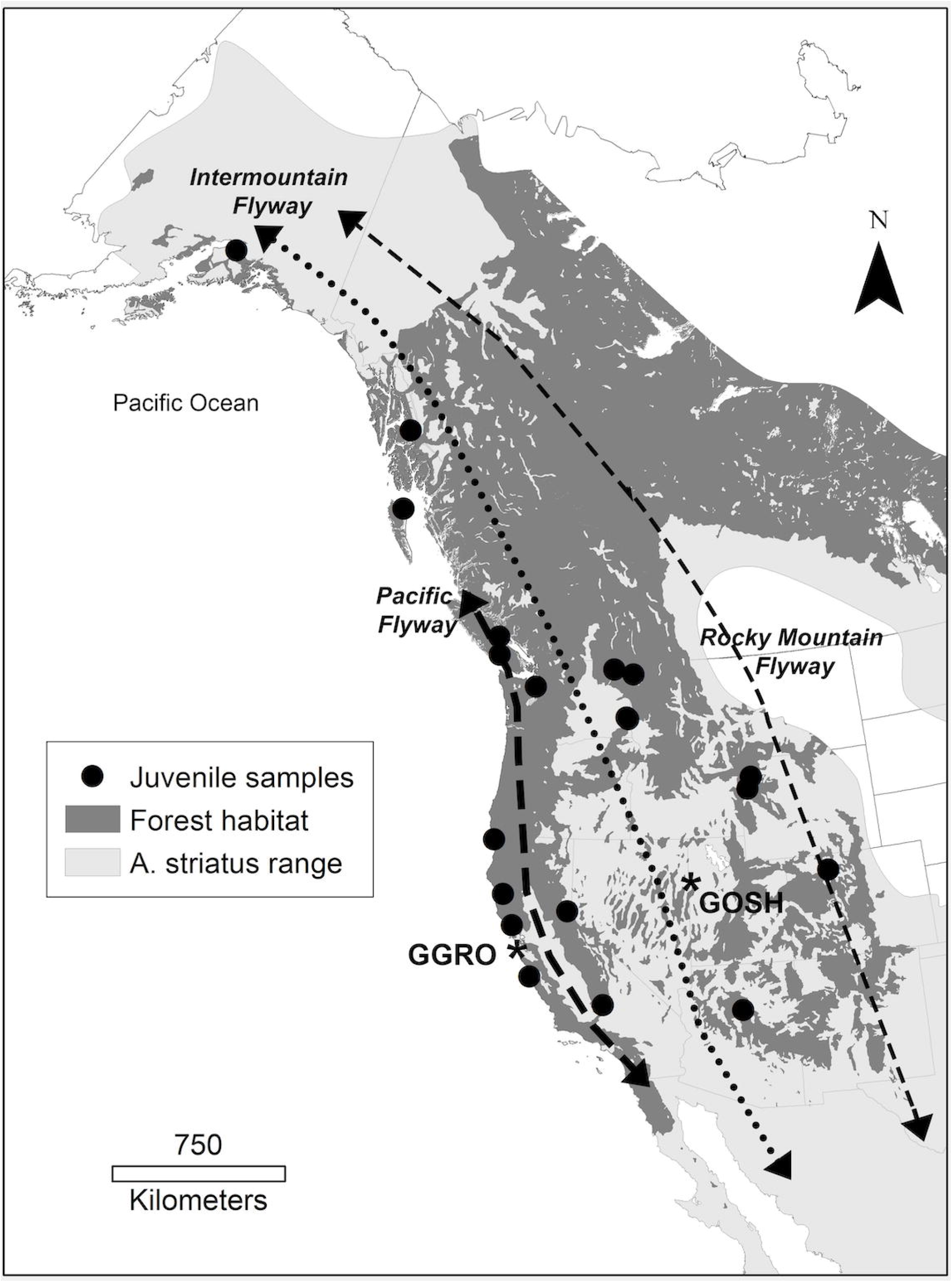
Map of sample locations for museum specimens of juvenile Sharp-shinned Hawk (*Accipiter striatus*) feathers. Museum specimens (*n* = 23) mapped in reference to the species known range in Western North America (light gray), and suitable forest habitat (dark gray). Juvenile samples (*n* = 23) are shown as triangles. Collection sites of migratory bird samples are indicated as GGRO (Golden Gate Raptor Observatory) and GOSH (Goshutes Mountains). The dominant migratory flyways of western North America are indicated by dashed lines and modified from Hoffman et al. (2002) for illustrative purposes. State and country boundaries are modified from public domain GIS files, US Census Bureau (2016) and Natural Earth (2020). Species range acquired with permission from BirdLife International and NatureServe (2015), and data to create the GIS biome layer acquired from Brown, Bennan, and Unmack (2007).

**Table 1.** Hydrogen and oxygen stable isotope composition of feathers of museum juvenile Sharp-shinned Hawk (*Accipiter striatus*) specimens (δ^2^H_f_ and δ^18^O_f_ values (‰)), and IsoMAP isoscape modeled stable hydrogen and oxygen isotope composition of precipitation (δ^2^H_p_ and δ^18^O_p_ values (‰)) of known natal origin.

| <sup>a</sup>Museum | <sup>b</sup>Specimen no. | Life stage | <sup>c</sup>State | Latitude | Longitude | $\delta^2\text{H}_f$ (‰) | $\delta^{18}\text{O}_f$ (‰) | $\delta^2\text{H}_p$ (‰) | $\delta^{18}\text{O}_p$ (‰) |
| --- | --- | --- | --- | --- | --- | --- | --- | --- | --- |
| **MVZ** | 169006 | Juvenile | AK | 61.0561111 | -149.79722 | -80.3 | 12.88 | -124.9 | -13.23 |
| **MVZ** | 99707 | Juvenile | BC | 54.0167 | -132.15 | -39.3 | 12.09 | -95.5 | -12.07 |
| **MVZ** | 32217 | Juvenile | CA | 36.537017 | -121.9262 | -18.7 | 16.23 | -36.9 | -4.36 |
| **MVZ** | 144621 | Juvenile | ID | 48.4797 | -116.8483 | -46.7 | 16.99 | -102.8 | -12.79 |
| **MVZ** | 15622 | Juvenile | BC | 48.9895 | -124.8 | -70.5 | 8.76 | -72.0 | -8.95 |
| **MVZ** | 81827 | Juvenile | BC | 49.6833 | -124.9333 | -44.0 | 13.03 | -71.9 | -10.02 |
| **MVZ** | 99705 | Juvenile | BC | 54.0167 | -132.15 | -44.9 | 12.69 | -95.5 | -12.07 |
| **MVZ** | 99706 | Juvenile | BC | 54.0167 | -132.15 | -58.3 | 13.07 | -95.5 | -12.07 |
| **MVZ** | 30835 | Juvenile | CA | 39.6863371 | -123.48519 | -12.1 | 22.22 | -51.2 | -5.65 |
| **MVZ** | 9775 | Juvenile | AK | 57.03139 | -132.8536 | -65.1 | 11.60 | -122.4 | -14.60 |
| **CAS** | 58055 | Juvenile | CA | 39.14057 | -120.2011 | -60.5 | 15.76 | -83.0 | -5.98 |
| **CAS** | 87247 | Juvenile | CA | 41.78846 | -124.1668 | -34.9 | 13.12 | -49.2 | -5.55 |
| **CAS** | 98830 | Juvenile | CA | 38.49657 | -122.9394 | -26.0 | 16.72 | -42.4 | -4.76 |
| **CRCM** | 57-367 | Juvenile | WA | 48.66223 | -117.9829 | -75.4 | 13.04 | -97.6 | -12.18 |
| **CRCM** | 57-368 | Juvenile | WA | 48.66223 | -117.9829 | -81.1 | 13.26 | -97.6 | -12.18 |
| **CRCM** | 89-229 | Juvenile | WA | 46.79025 | -117.2521 | -87.0 | 10.05 | -94.2 | -9.72 |
| **CRCM** | 89-223 | Juvenile | WA | 46.73064 | -117.1625 | -67.4 | 11.41 | -94.2 | -10.01 |
| **UWYMV** | 1352 | Juvenile | WY | 44.27615 | -110.4736 | -103.7 | 8.62 | -102.5 | -11.99 |
| **UWYMV** | 2521 | Juvenile | WY | 43.83333 | -110.7 | -75.1 | 13.41 | -100.0 | -11.47 |
| **UWYMV** | 2547 | Juvenile | CO | 40.39162 | -106.9051 | -62.2 | 15.97 | -82.7 | -9.67 |
| **SDNHM** | 52830 | Juvenile | WA | 47.8674 | -122.516 | -35.6 | 10.90 | -69.0 | -8.40 |
| **SDNHM** | 54234 | Juvenile | AZ | 35.1957 | -111.6326 | -26.0 | 17.19 | -52.0 | -5.78 |
| **SDNHM** | 18938 | Juvenile | CA | 35.50708 | -118.3434 | -85.3 | 10.08 | -64.8 | -4.72 |

<sup>a</sup>Museums: CAS = California Academy of Sciences, San Francisco, CA, USA; CRCM = Charles R. Conner Museum, Pullman, WA, USA; MVZ = Museum of Vertebrate Zoology, University of California, Berkeley, CA, USA; SDNHM = San Diego Natural History Museum, San Diego, CA, USA; and UWYMV = University of Wyoming Museum of Vertebrates, Laramie, WY, USA.
<sup>b</sup>Specimens: Full information on each specimen can be obtained by taking the specimen number and searching for it in the online databases in http://vertnet.org/index.html and https://arctosdb.org/
°State or Provinces: AK = Alaska, USA; AZ = Arizona, USA; BC = British Colombia, Canada; CA = California, USA; CO = Colorado, USA; ID = Idaho, USA; NM = New Mexico, USA; NV = Nevada, USA; OR = Oregon, USA; UT = Utah, USA; WA = Washington, USA; WY = Wyoming, USA.

### Sample preparation and SIA

Feathers were cleaned by immersion in a chloroform:methanol 2:1 (v/v) solution for 24 hours, and then again for 1 hour, to remove lipids and mites. After each wash the feathers were air dried for 24 hours [58–59]. The barbs were removed from the rachis, minced, and 1.5-2.0 mg of feather material was packed into 3.5 x 5 mm silver capsules. To avoid the possibility of intrafeather variation found in distal versus proximal samples for some raptor feathers, barbs for oxygen and hydrogen samples were removed from corresponding sides of each contour feather [57]. Previous work on other species of raptors has found that breeding condition and age may affect δ^2^H_F_ values differently in juvenile versus adults [60–62]. Therefore, adult samples were removed from analysis reflecting these concerns. Only samples from juvenile individuals were analyzed for both δ^18^O_F_ and δ^2^H_F_ values (museum: *n* = 23, migratory: *n* = 21). However, as there are no published results of the differences for stable oxygen isotope measurements between adult and juvenile birds of prey, we present the δ^18^O values for adult *A. striatus* samples of know origin (*n* = 25) as Supporting Information in this paper (S1 Table, S1 Fig).

The stable isotope abundances are presented in δ notation as deviations from the standard reference (V-SMOW) in parts per mil (‰) according to the following equation: δ X = (R_sample_/R_standard_) −1) where X represents ^2^H or ^18^O, and R the ratio of the heavy and light isotope (e.g., ^18^O/^16^O) in the sample and in the standard, respectively.

The δ^18^O measurements were performed at the Center for Stable Isotope Biogeochemistry (University of California, Berkeley, CA, USA) using a PYRO Cube (Elementar, Hanau, Germany) interfaced to a Thermo Delta V mass spectrometer (Thermo Fisher Scientific Inc., Waltham, Massachusetts, USA). The δ^2^H measurements were carried out at the Cornell Isotope Laboratory (Cornell University, Ithaca, NY, USA) using a Temperature Conversion Elemental Analyzer (TC/EA) interfaced to a Thermo Delta V mass spectrometer (both from Thermo Fisher Scientific Inc., Waltham, Massachusetts, USA). Both measurements were based on pyrolysis of the sample carried out in reactors kept at 1350 °C, with GC temperature kept at 90°C for hydrogen analysis, and CO trap temperature kept at 40 °C and then increased to 130 °C for release of CO for oxygen analysis.

Reference materials of kudu horn and caribou hoof keratins (KHS and CBS) were used for normalization (δ^18^O value = +20.3 ‰ and +3.8‰, respectively [63]; δ^2^H value = −35.3 ‰ and −157.0 ‰, respectively [64]), and an in-house keratin material was used as quality control material for both hydrogen and oxygen stable isotope measurements. The precision of the analysis was ± 2.8‰ and 0.30‰ for hydrogen and oxygen, respectively. To correct the measured hydrogen isotope ratios for the contribution of exchangeable hydrogen atoms to the total number of hydrogen atoms in feathers (~15%) [47, 63, 65], samples were allowed to equilibrate with the laboratory ambient atmosphere along with KHS and CBS standards for a minimum of 72 hours before isotopic analysis [65].

A linear regression approach was used to examine the relationship between measured δ^2^H_F_ and δ^18^O_F_ values from *A. striatus* feathers for museum samples of known natal/breeding origin as well as in birds sampled at the two migratory banding sites.

### H and O precipitation Isoscapes for *Accipiter striatus* museum samples of known natal/breeding origin

IsoMAP, a web resource for isoscapes modeling, (http://www.waterisotopes.org)[66], was used to predict the hydrogen and oxygen isotopic compositions of precipitation (δ^2^H_P_ and δ^18^O_P_ values) at the locations where museum specimens were collected. Isoscapes are gridded surfaces representing spatially explicit isotope distributions across a landscape [46, 67]. IsoMAP software creates water isoscapes by using precipitation isotope data from the Global Network for Isotopes in Precipitation (GNIP) database administered by the International Atomic Energy Association and World Meteorological Organization [68]. Within IsoMAP, numerous parameters can be selected to create an isoscape model that is the best fit for a specific study area and time period since the geographical variation of δ^2^H_P_ and δ^18^O_P_ values depend on a range of geographic and meteorological effects including latitude, season, elevation, and regional air mass circulation. Independently for each isotope, we modeled the geographic distribution of δ^2^H_P_ and δ^18^O_P_ values across North America using precipitation data collected from 1960–1999 during the months from March to September to represent the plant growing season [68, 69]. Comparisons of different precipitation isotope models available in IsoMAP revealed that the most robust isoscapes that represented δ^2^H_P_ and δ^18^O_P_ values across North America were kriging interpolation models. The δ^2^H_P_ isoscape produced was based on 117 stations, had resolution of 9×9 km, a correlation parameter of 0.93, and included the variables elevation (ETOPO, *P* < 0.001), latitude (*P* < 0.001) and longitude (*P* = 0.06) (available as IsoMAP job key 50333, S2 Fig [70]). The most robust δ^18^O_P_ isoscape was based on 120 stations, had resolution of 9×9 km, a correlation parameter of 0.92, and included the variables elevation (ETOPO, *P* < 0.001), latitude (*P* < 0.001) and longitude (*P* = 0.05) (available as IsoMAP job key 63026, S2 Fig [71]).

The hydrogen and oxygen isoscape precipitation models were further modified using the spatial software ArcGIS (ESRI 2010). In particular, we limited the spatial isotopic predictions to the known habitat range of *A. striatus* using a geographic information system (GIS) layer of range delineation provided by BirdLife International and NatureServe [72]. In addition, because this species is found to nest specifically in forests [21], we applied a GIS biome layer to exclude non-breeding habitat, such as tundra, open water, desert, and grassland from potential sites of origin [73].

### Predicting the origin of migrating *Accipiter striatus*

Because of isotope discrimination during feather formation and other effects, the isotope values of feathers may not directly reflect the stable hydrogen and oxygen isotopic composition of the environmental water at the site where they are formed [74–75]. We estimated the magnitude of such discrimination factors for *A. striatus* feathers by calculating the linear relationship between the isotopic compositions of feathers from museum specimens of known origin (δ^2^H_F_ and δ^18^O_F_) and the corresponding isoscape predicted precipitation values (δ^2^H_P_ and δ^18^O_P_) at the location where the museum specimens were collected [76–78]. Metadata associated with each specimen included the uncertainty for the collection location in meters. When multiple isoscape derived δ^2^H_P_ and δ^18^O_P_ grid values were available within the radius of uncertainty around a collection location, we used the average isotope values within this radius.

The parameters of the linear regression equations derived from the museum specimens data were used to predict the isotopic composition of the precipitation at the site of origin of birds sampled at each of the two migratory banding sites (GGRO and Goshutes). The feather isotopic compositions of the migrating birds represented the variable *x* while *y* represented an estimate of the associated δ^2^H_P_ or δ^18^O_P_ values. Resulting δ^2^H_P_ and δ^18^O_P_ values thus represented the water source isotope compositions expected for the localities where the feathers of migrating birds were formed. The linear regressions were used as a transfer function to convert feather isotope values for migratory birds to precipitation isotopes values at the sites of origin [79]. Many researchers examining the relationship between feather and precipitation isotopes plot precipitation on the x axis and feather isotopes on the y axis as the dependent variable, creating a feather isoscape [39, 40, 48, 51]. However, we did not feel justified using this format of regression equation for the transformation lines in our analysis with the sample size of museum specimens (*n* = 23) across the entire area of interest. Instead we predicted precipitation from measured feather values and created a precipitation isoscape to predict probability of origin. For comparison with previous analyses from the literature we have created the regression equations in the supporting information (S4 Fig).

To determine if the origin for birds migrating through the GGRO and Goshutes sites differed, we examined the data in several ways. First, we calculated the frequency distributions of the predicted precipitation isotope values for the two migratory banding sites. Second, δ^2^H_F_ and δ^18^O_F_ values for each migrant group (GGRO and Goshutes) were compared statistically using Welches unequal variance T-test which tests the hypothesis that two populations have the same means. Additionally, δ^2^H_F_ and δ^18^O_F_ values for each migrant group (GGRO and Goshutes) were examined with a Kolmogorov-Smirnov distribution test which tests whether the two populations have the same distribution.

Finally, we utilized the IsoMAP geographic assignment function to produce maps representing the likelihood of origin for the migrant birds sampled at GGRO and Goshutes. The assignment function in IsoMAP uses a semi-parametric Bayesian framework to model probability density surfaces that can be used to determine geographic areas where organic material, such as bird feathers, were developed [79, 80]. The assignment function requires an observed sample isotopic composition, a standard deviation associated with the environment to sample transfer, and an isoscape model in IsoMAP that the sample is compared to. For each of the birds sampled at the GGRO and Goshutes, δ^2^H_P_ and δ^18^O_P_ values were estimated using the feather to precipitation linear regressions. The residual standard error (RSE) of each linear regression was used as the estimate of error (RSE for δ^2^H of 18.3‰, and RSE for δ^18^O of 3.0). The precipitation isoscape models described above (IsoMAP job key 50333 and 63026, S2 Fig) [70, 71] were utilized in the assignment function. Additional uncertainty associated with the precipitation isoscape is automatically included in the assignment algorithm.

Rather than creating average probability surfaces for each migratory group though the bulk sample function, probability surfaces were generated for each individual bird [80]. The individual probability density maps were then averaged for GGRO and Goshutes groups. Using ArcGIS, individual rasters within each group were summed together and then normalized by the sum of all cell values in the final density surface. This process resulted in geographic representations of the likely origin of *A. striatus* migrating along the Pacific Coast and the Intermountain Flyways. Areas outside the species range and nesting habitat type were not included as likely origin areas of the migratory birds. To examine the accuracy of the linear transfer functions and the assignment of origin models, we also generated probability density surfaces for 10 museum samples of known origin (S5 Fig, Table S2). All statistics were performed in R (R version 2.14.0) [81].

## Results

The stable hydrogen and oxygen isotopic composition of feathers of juvenile *A. striatus* museum specimens varied from −103.7 to −12.1 ‰ for δ^2^H_F_ and from 8.62 to 22.22 ‰ for δ^18^O_F_ respectively (*n* = 23) (Table 1). A significant and positive relationship was found between the δ^2^H_F_ and δ^18^O_F_ values from the juvenile feathers (R^2^ = 0.48, *P* < 0.001, y = 5.34(x) – 128.95) (Fig 2).

**Fig 2.**
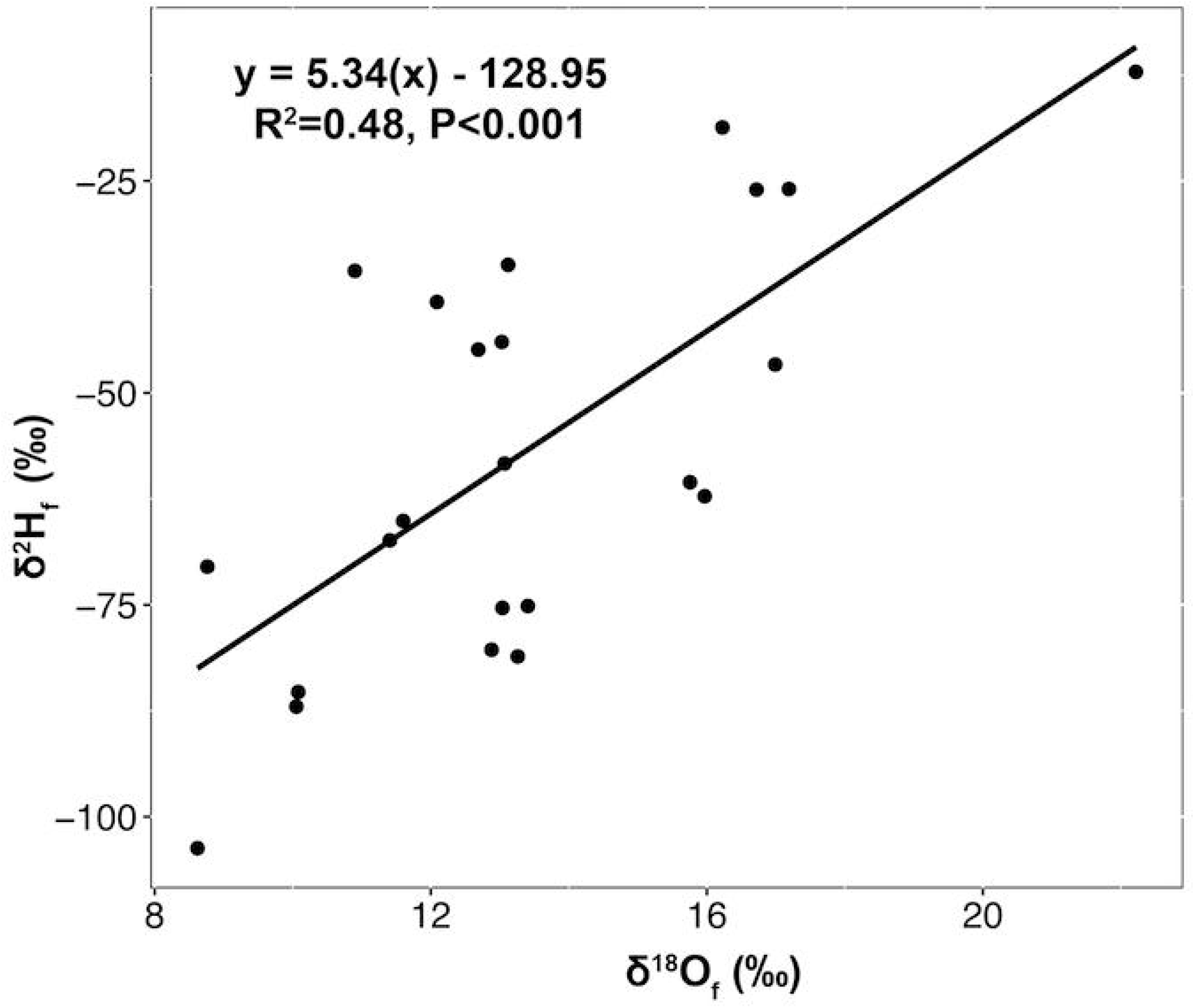
Relationship between the stable isotope composition of museum specimens of Sharp-shinned Hawks (*Accipiter striatus*). Stable hydrogen (δ^2^H_F_ ‰) and oxygen (δ^18^O_F_ ‰) isotope values for juvenile museum feather specimens (*n* = 23) of known natal origin.

We found positive relationships between the feather isotope values and the isoscape modeled precipitation isotope values for the *A. striatus* museum specimen of known origin. The linear regression between δ^2^H_F_ and δ^2^H_P_ values was statistically significant (*n* = 23, R^2^ = 0.46, *P* < 0.001, y = 0.68x – 43.98, RSE = 18.3‰) (Fig 3a), while the linear regressions between δ^18^O_F_ and δ^18^O_P_ values based on feathers of juvenile birds (Fig 3b) were not statistically significant (*n* = 23, R^2^ = 0.14, *P* = 0.07, *y* = 0.39*x* – 14.67, RSE = 3.0‰).

**Fig 3.**
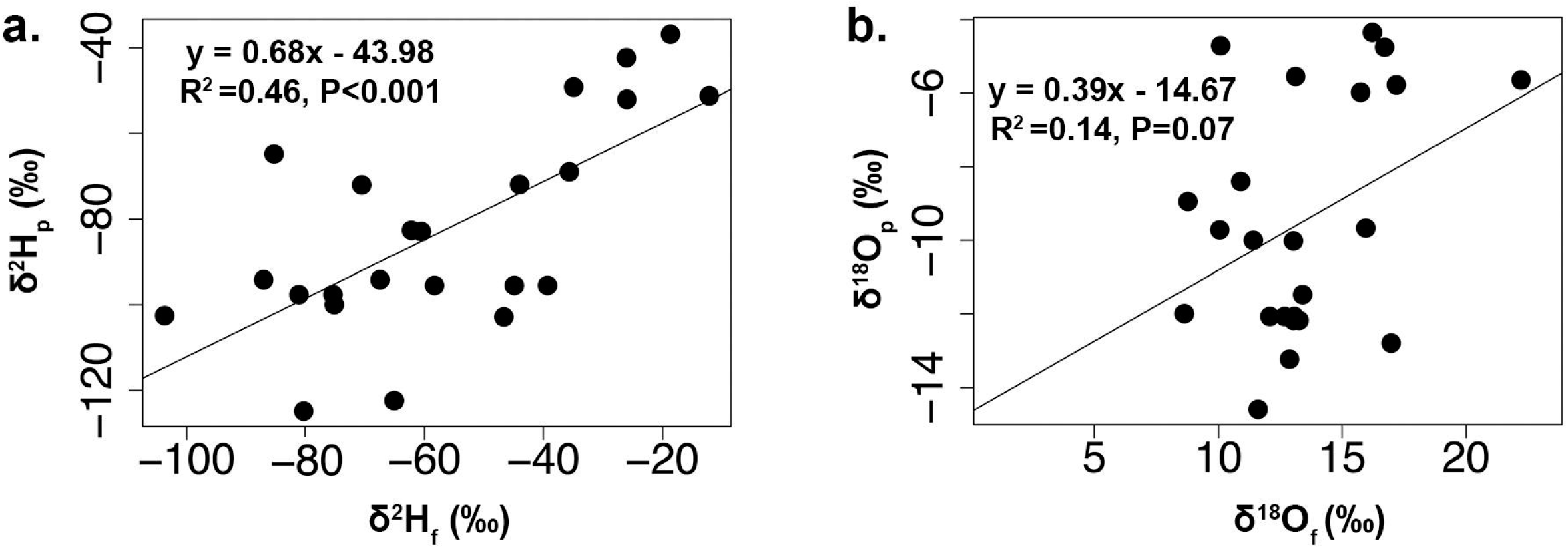
Relationship between the stable isotopic compositions of museum feathers and the isoscape model of precipitation. Stable hydrogen (δ^2^H_F_ ‰) and oxygen (δ^18^O_F_ ‰) isotopic composition of feathers for museum Sharp-shinned Hawk (*Accipiter striatus*) specimens of known natal origin and the isoscape modeled isotopic compositions of precipitation (δ^2^H_P_ and δ^18^O_P_ ‰) at the collection locations: (a) δ^2^H_F_ values of birds (*n* = 23) versus δ^2^H_P_ values, (b) δ^18^O_F_ values of birds (*n* = 23) versus δ^18^O_P_ values.

Within the set of migratory birds for which both hydrogen and oxygen isotopic compositions were measured, there was a positive relationship between δ^2^H_F_ and δ^18^O_F_ values at both the GGRO and Goshutes banding sites (Fig 4). The linear relationship between δ^2^H_F_ and δ^18^O_F_ values for the birds captured at the GGRO was very similar to that found for the museum bird feathers (*n* = 14, R^2^ = 0.42, *P* = 0.01, y = 6.36x – 146.45, Fig 4a). However, the relationship between δ^2^H_F_ and δ^18^O_F_ values for the birds migrating through the Goshutes banding site was not statistically significant (*n* = 7, R^2^ = 0.18, *P* = 0.34, y = 1.02x – 122.29, Fig 4b).

**Fig 4.**
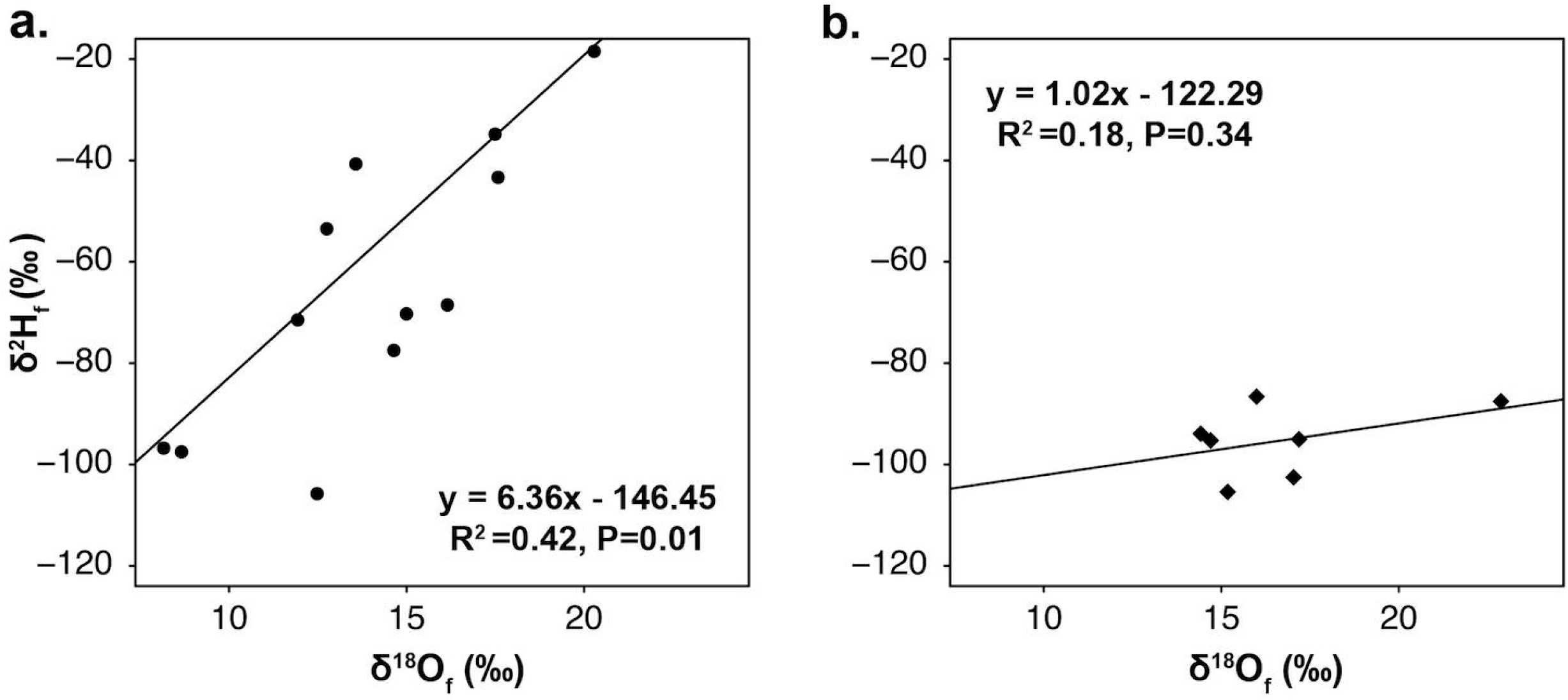
Relationships between the stable isotope compositions of feathers from birds trapped along the migratory flyways. Stable oxygen (δ^18^O_F_ ‰) and hydrogen (δ^2^H_F_ ‰) isotope composition of feathers for juvenile migratory Sharp-shinned Hawk (*Accipiter striatus*) specimens collected (a) along the Pacific Coast Flyway at the migratory banding site the Golden Gate Raptor Observatory (GGRO, *n* = 14) and (b) along the Intermountain Flyway at the migratory banding site Goshute Mountains HawkWatch (Goshutes, *n* = 7).

In migratory bird feathers, the range of δ^2^H_F_ values was −105.69 to −4.36 ‰ from the GGRO (*n* = 15) and −105.40 to −86.60 ‰ at the Goshutes (*n* = 7) (Figure 5a and c, Table 2). The variation in δ^18^O_F_ values was 8.16 to 21.19 ‰ for the GGRO (*n* = 19) and 14.11 to 22.87 ‰ for the Goshutes (*n* = 10) (Figure 5b and d, Table 2). The mean δ^2^H_F_ values (−58.6 ± 32.7 ‰ GGRO, −95.7 ± 7 ‰ Goshutes) were statistically different between the GGRO and Goshutes sites (Welch’s two-sample t-test, t = 4.13, df = 16.5, *P*< 0.001). There was a significant difference in the distribution of the δ^2^H_F_ values for the two migratory flyways (Two-sample Kolmogorov-Smirnov test, D = 0.73, *P* = 0.005). Although a subsample of birds captured at the GGRO site showed δ^18^O_F_ depleted values compared to the Goshutes birds, the mean δ^18^O_F_ values (14.78 ± 3.5 GGRO, 16.44 ± 2.7 Goshutes) did not differ statistically between sites (Welch’s two-sample t-test, t = −1.4, df = 23.2, *P* = 0.16). Also, we did not detect any significant difference in the distribution of the δ^18^O_F_ values for the migratory flyways (Two-sample Kolmogorov-Smirnov test, D = 0.42, *P* = 0.16).

**Fig 5.**
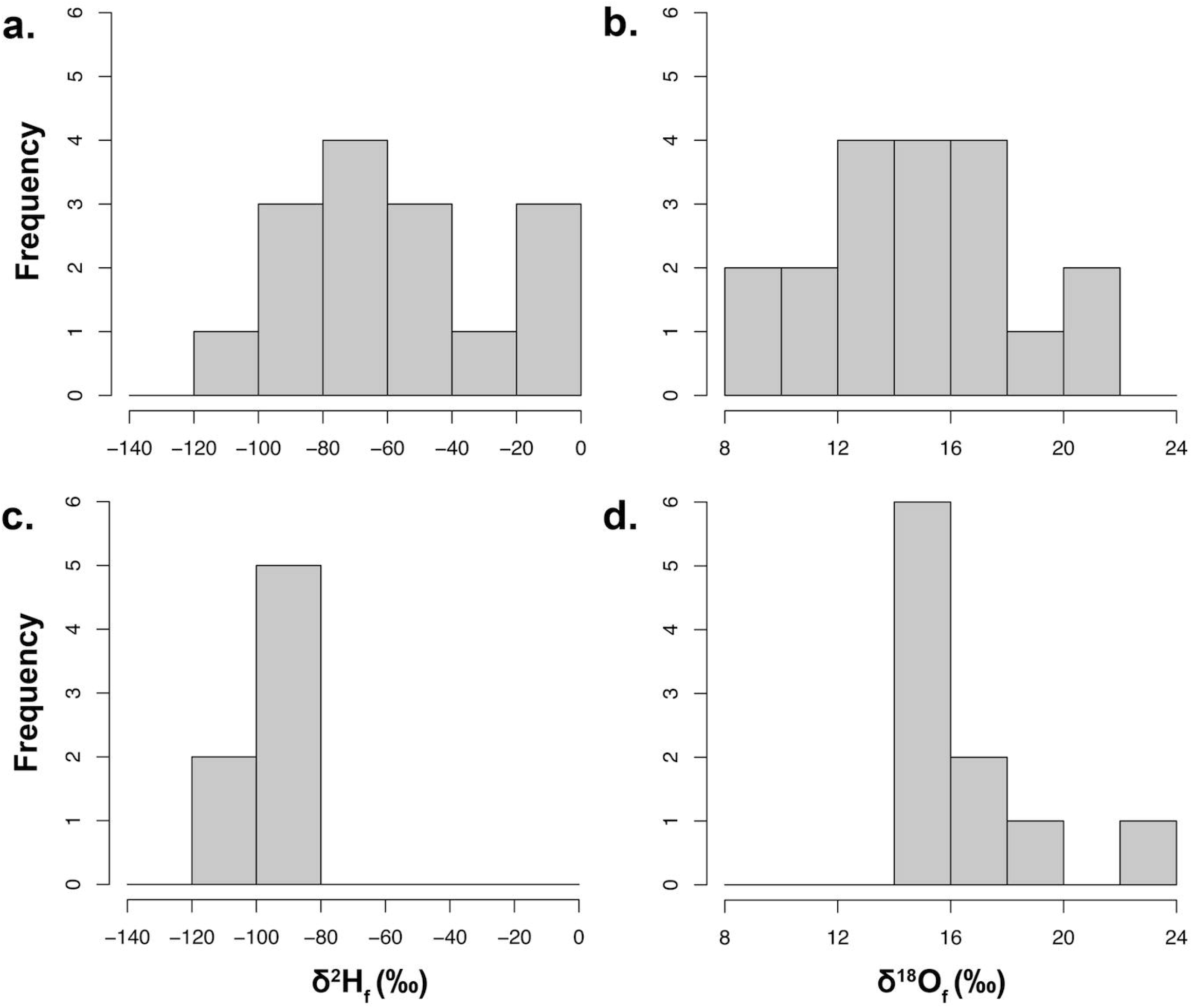
Frequency distribution of the isoscape modeled isotope compositions of feathers of migratory Sharp-shinned Hawks. Predicted stable hydrogen (δ^2^H_F_ ‰) and oxygen (δ^18^O_F_ ‰) isotopic compositions of precipitation at the natal origin for migratory Sharp-shinned Hawk (*Accipiter striatus*) specimens collected (a and b) along the Pacific Coast Flyway at the Golden Gate Raptor Observatory (GGRO, *n* = 17), and (c and d) along the Intermountain Flyway at the Goshute Mountains HawkWatch (Goshutes, *n* = 10). Values on the y-axis represent counts of individual specimens.

**Table 2.** Stable hydrogen and oxygen isotope composition of feathers of migrating Sharp-shinned Hawks (*Accipiter striatus*) (δ^2^H_f_, and δ^18^O_f_ values (‰)) captured at the Golden Gate Raptor Observatory (GGRO), and at the Goshute Mountains HawkWatch (Goshutes,), and of predicted stable hydrogen and oxygen isotope composition of precipitation (δ^2^H_p_ and δ^18^O_p_ values (‰)) at the migrant’s unknown natal origin.

| Migratory Banding site | <sup>a</sup>Sample Band number | <sup>b</sup>$\delta^2\text{H}_f$ (‰) | <sup>b</sup>$\delta^{18}\text{O}_f$ (‰) | <sup>b, c</sup>$\delta^2\text{H}_p$ (‰) | <sup>b, c</sup>$\delta^{18}\text{O}_p$ (‰) |
| --- | --- | --- | --- | --- | --- |
| GGRO | 1363-74740 | -70.20 | 14.99 | -91.74 | -8.90 |
|  | 1353-16673 | -68.60 | 16.16 | -90.60 | -8.45 |
|  | 1162-13236 | -18.50 | 20.28 | -56.58 | -6.86 |
|  | 733-23478 | -53.50 | 12.76 | -80.37 | -9.76 |
|  | 1363-74746 | -40.70 | 13.58 | -71.67 | -9.44 |
|  | 2003-95304 | -77.50 | 14.65 | -96.66 | -9.03 |
|  | 0733-64092 | -105.69 | 12.47 | -115.69 | -9.87 |
|  | 1423-50079 | -71.40 | 11.93 | -92.54 | -10.08 |
|  | 1433-84282 | -97.44 | 8.66 | -110.24 | -11.33 |
|  | 1423-50062 | -43.30 | 17.57 | -73.44 | -7.90 |
|  | 2003-95657 | -34.73 | 17.49 | -67.60 | -7.94 |
|  | 1433-84273 | -9.04 | 13.42 | -50.13 | -9.50 |
|  | 2003-95656 | -96.80 | 8.16 | -109.80 | -11.53 |
|  | 2003-95572 | -4.36 | 15.67 | -46.94 | -8.64 |
|  | 2003-95363 | -87.90 |  | -103.75 |  |
|  | 1152-29272 |  | 16.11 |  | -8.47 |
|  | 2003-95575 |  | 21.19 |  | -6.51 |
|  | 1363-74737 |  | 18.86 |  | -7.41 |
|  | 2003-95653 |  | 15.13 |  | -8.85 |
|  | 1152-29376 |  | 11.66 |  | -10.18 |
| Goshutes | 1523-88735 | -102.50 | 17.04 | -113.70 | -8.11 |
|  | 1162-71742 | -95.30 | 14.70 | -108.75 | -9.01 |
|  | 1523-88624 | -105.40 | 15.18 | -115.65 | -8.83 |
|  | 1523-88749 | -86.6 | 15.99 | -102.87 | -8.51 |
|  | 1162-71901 | -87.50 | 22.87 | -103.51 | -5.86 |
|  | 1523-88752 | -95.00 | 17.19 | -108.59 | -8.05 |
|  | 1523-88734 | -93.90 | 14.42 | -107.84 | -9.12 |
|  | 1162-71551 |  | 14.11 |  | -9.24 |
|  | 1523-88628 |  | 18.36 |  | -7.60 |
|  | 1162-71547 |  | 14.58 |  | -9.06 |

<sup>a</sup> Band numbers represent individual numbers issued by the Bird Banding Laboratory
(http://www.pwrc.usgs.gov/BBL/bblretrv/)
<sup>b</sup> Missing values represent feathers where there was not enough sample from one specimen for both hydrogen and oxygen analysis.
<sup>c</sup> The $\delta^2\text{H}_p$ values were predicted using the linear regression equation $\delta^2\text{H}_p = 0.68*\delta^2\text{H}_f - 43.98$ (RSE = 18.3‰).
The $\delta^{18}\text{O}_p$ values were predicted using the linear regression equation $\delta^{18}\text{O}_p = 0.39*\delta^{18}\text{O}_f - 14.67$ (RSE = 3.0‰).

The predicted δ^2^H_P_ values for migratory birds from the GGRO banding site varied between −46.94 and −115.69 ‰ (Table 2). The average of these probability density surfaces shows that the origin of the individuals captured along the Pacific Coast Flyway are most likely from eastern CA, Oregon (OR), and Washington (WA) as well as some forested areas within NV, Utah (UT), Colorado (CO), Montana (MT), Idaho (ID), Wyoming (WY), British Columbia (Canada), and southern Alaska (AK) (Fig 6a). For the birds that migrated through the Goshutes banding site, the predicted δ^2^H_P_ values were less varied than at the GGRO, ranging between −102.87 and −115.65‰ (Table 2). The average probability density surface suggests that Intermountain Flyway captured birds originated from a smaller area focused primarily in eastern WA, WY, MT, ID, British Columbia (Canada), and in southern AK (Fig 6c). The δ^2^H_P_ probability surfaces generated for museum samples of known origin showed a high to medium probability of prediction for the correct localities for almost all the specimens, except for a specimen from the western slope of the Rocky Mountains in ID (S5 Fig).

**Fig 6.**
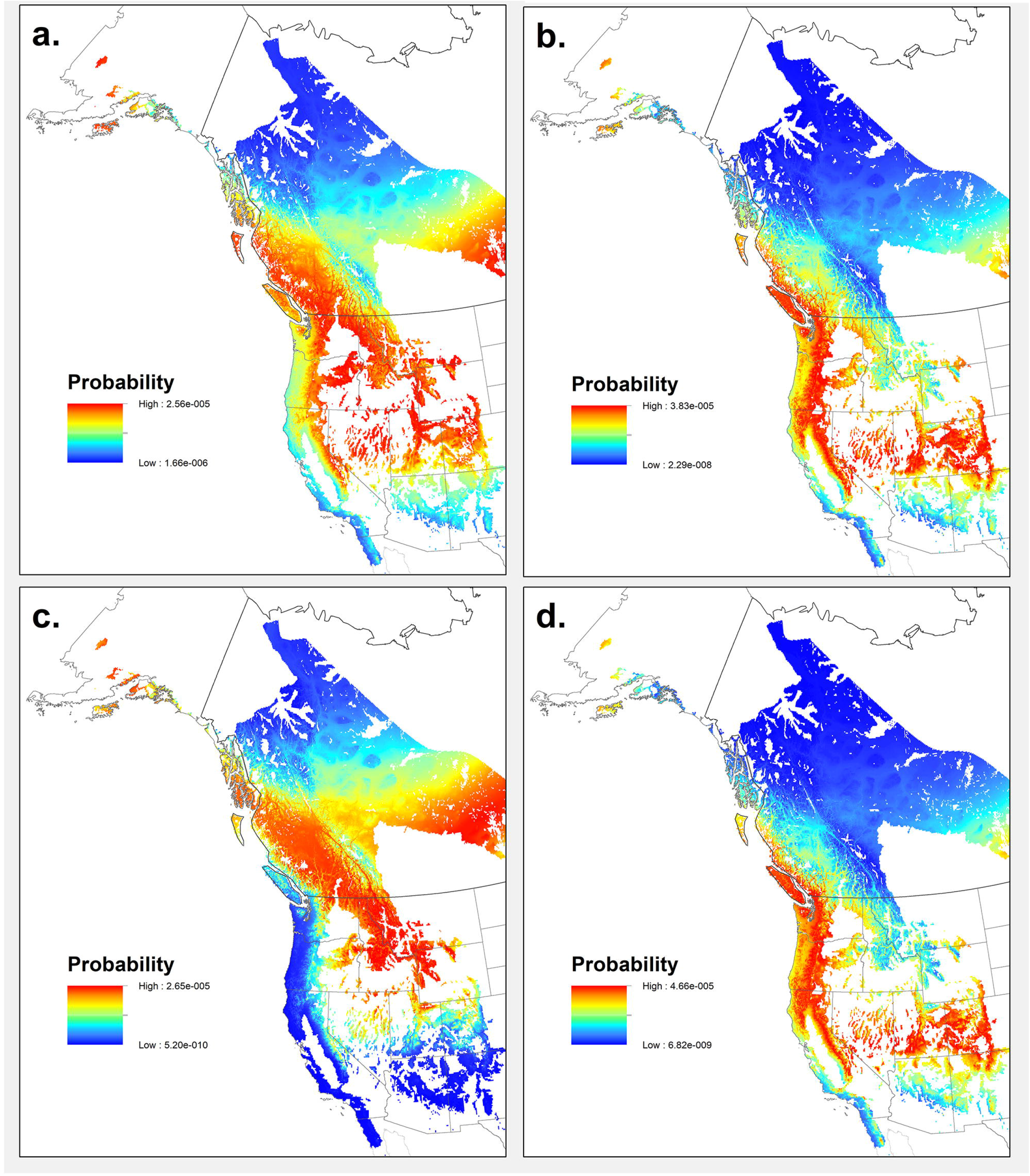
Probability density maps of the origin of migrating juvenile Sharp-shinned Hawks (*Accipiter striatus*). Maps are based on predicted δ^2^H_P_ values (‰) (left panels) and predicted δ^18^O_P_ values (‰) (right panels) for birds captured (a and b) along the Pacific Coast Flyway at the Golden Gate Raptor Observatory (GGRO) and (c and d) along the Intermountain Flyway at the Goshute Mountains HawkWatch (Goshutes). Each map represents the mean of probability density surfaces created for individual birds sampled at a location and by isotope group. State and country boundaries are from public domain GIS files US Census Bureau (2016) and Natural Earth (2020). Species range acquired with permission from BirdLife International and NatureServe (2015), and data to create the GIS biome layer acquired from Brown, Bennan, and Unmack (2007).

Isoscape predicted δ^18^O_P_ values at the natal/breeding sites of the birds captured at the GGRO site ranged between −6.51 and −11.53‰ (Table 2). The average probability density surface for birds captured along the Pacific Coast Flyway showed that the natal origins were most likely from eastern CA, OR, NV, UT, CO, MT, WA, coastal British Columbia (Canada) and southern AK (Fig 6b). These predicted origins, in general, confirmed the data found from the δ^2^H analysis. Predicted δ^18^O_P_ values at the origin sites of the birds captured at the Goshutes ranged from −5.86 to −9.24‰ (Table 2). The average probability density surface based on oxygen isotope composition located the natal origins of birds from the Intermountain Flyway in more coastal and southern areas then those from the δ^2^H analysis, including CA, NV, UT, CO, OR, WA, British Columbia (Canada), and small portions of AK (Fig 6d). Similar to hydrogen the δ^18^O_P_ probability density surfaces generated for museum samples of known origin showed high to medium to medium probability of prediction for most specimens, but showed low probability of origin for two specimens collected along the western slope of the Rocky Mountains in ID and WA, and a specimen collected from the southern coast of AK (S5 Fig).

## Discussion

Results from the combined SIA of hydrogen and oxygen of *A. striatus* feathers showed that some raptors migrating along the Pacific Coast Flyway have origins that overlap with those of raptors migrating along the Intermountain Flyway. Our prediction that juvenile birds that traveled along each migratory route would come from different and non-overlapping breeding/natal origins was therefore not supported. Instead it appears that migratory *A. striatus* juveniles, specifically those that travel along the Pacific Coast Flyway may come from both west of the Sierra Nevada mountain range and from the northern Rocky Mountain Range and western interior regions of North America (Fig 6).

This outcome should be taken into consideration for conservation of *A. striatus* in western North America, as individual populations may not show specific adaptations or fidelity for movement along a single migratory flyway [7, 10, 11]. Instead, adjustments to their migration strategy may depend on a multitude of ecological factors, such as minimization of energy cost and mortality risk [82]. Juvenile raptors have also been found to have a greater degree of variation in their migratory movement then adults [83], perhaps leading juvenile *A. striatus* to wander between flyways during their first migration. As a result, movement across multiple flyways should be heeded when examining data from migratory watch and banding sites, and when using this data to estimate population size and fluctuations.

In agreement with previous studies, we found that both stable hydrogen and oxygen isotopic composition of feathers can be used to predict the origin of birds across broad spatial scales [39, 40]. However, the methodology based on oxygen had, in general, less predictive power than the one based on hydrogen. Within the overall findings of natal origins for the birds migrating along western migratory flyways in North America, some differences were found in the predicted natal sites depending on which isotope and migratory route were examined.

### Assignment of origin

We found an overlap in natal origin of *A. striatus* for the Flyways based on hydrogen and oxygen isoscapes (Fig 6). Birds migrating along the Pacific Coast Flyway originated from a larger area that covered both the coastal and central areas of the species’ western range while *A. striatus* migrating along the Intermountain Flyway had a more limited geographical origin.

Results from both hydrogen and oxygen isotopes point to similar predicted natal areas for *A. striatus* migrating along the Pacific Coast Flyway (Fig 6a and b) in forest sites in CA, NV, OR, WA, ID, and British Columbia (Canada). The origins of the migrants from the Intermountain Flyway predicted from the oxygen isoscape also included forests in the Sierra Nevada and Cascade mountains, spanning through CA, OR, and WA up into British Columbia (Fig 6d). This expands the natal territory of the migratory individuals from the Intermountain Flyway westward from the hypothesized origins, placing them closer to the west coast of North America, and creates substantial overlap with the natal origins of those from the Pacific Coast Flyway. In contrast, the origin sites for the Intermountain Flyway indicated using the hydrogen isoscape lie along our predicted areas of natal origin, and agree with previously published hydrogen data from Lott and Smith [51], which identified forests in the states of ID, MT, and British Columbia as origins for *A. striatus* migrating through this flyway (Fig 6c). As a result, the hydrogen isoscape revealed overlap in the origins of the two flyways only in the central areas of the species western range. Modeled high probability of origin for *A. striatus* from δ^2^H data was also found for regions associated with the central migration flyway, including the central Canadian providences of Alberta and Saskatchewan for both Flyways; but it is unclear if these values represent a true signal of origin or the lack of differentiation of δ^2^H_P_ values found in the central northern plains of North America [84]. The origin sites predicted using the oxygen isoscape data do not show the same high probability for either Flyway. An examination of the predicted origin of known museum samples did show low probability of origin for some specimens collected in ID and eastern WA for oxygen, suggesting that there may be bias in the assignment of origin from δ^18^O data for that area (S5 Fig). The weak relationship between the migrant’s δ^2^H_F_ and δ^18^O_F_ values may account for these discrepancy (Fig 4). However, weak correlations between δ^2^H_F_ and δ^18^O_F_ have not hindered previous studies that looked at the origin of birds in Europe and Asia [40], and the assignment of origin tests showed high probability for museum specimens with both hydrogen and oxygen in other areas of the species western range.

The successful use of SIA of hydrogen and oxygen to assign sites of origin for migratory animals depends on understanding the relationship between isotopic composition in tissues and the isotopic composition of waters sources within the landscape where these tissues were formed. Isotopic composition of animal tissues are offset from environmental isotope values due to a variety of discrimination factors that differ for each element [85]. These discrimination processes vary among specific isotopes. In this study, the relationship between the stable hydrogen and oxygen isotope composition of feathers differed depending on sampling locality. This relationship was stronger for both museum specimens of known origin (R^2^ = 0.48) and birds migrating through the Pacific Coast Migratory Flyways (R^2^ = 0.42) (Fig 2 and Fig 4a), than for birds traveling along the Intermountain Flyway (R^2^ = 0.18) (Fig 4b). Research on other species of vertebrates that compared δ^18^O and δ^2^H values in feathers, claws, or hair have also found varied correlation patterns between the two isotopic compositions. Significant correlations between δ^18^O and δ^2^H values were found for insectivorous passerines (R^2^ = 0.34) [39] and falcons (R^2^ = 0.64 and R^2^ = 0.48)[86, 48], as well as for herbivorous mammals (R^2^ = 0.84, and 0.57) [37]. However, no significant correlations were found between tissue oxygen and hydrogen isotope compositions for other vertebrate species, including Pumas (*Puma concolor*) [37] and European Cranes (*Grus grus*) [40]. Our study is the first to report that birds of the same species sampled along different migrating routes can show different relationships between δ^2^H_F_ and δ^18^O_F_ values.

The small sample size of birds captured along the Intermountain Flyways may also account for the poor relationship between the stable hydrogen and oxygen isotope composition in feathers. These samples were restricted to a single year of migratory data to coincide with sampling strategy of published genetic data and ensure that all the birds sampled belonged to the previously characterized *A. striatus* western population [53]. It may be possible that the individuals analyzed here represented a divergence from the average value for the site that could have been detected with a larger sample size [48]. However, the predicted origins from our assignments based on hydrogen isotopes align very closely with previously published work for *A. striatus* from the same flyway also based on hydrogen isotopes [51]. This suggests that despite a small sample size, our results agree with previous findings of origin of birds from the Intermountain Flyway. In general, increasing the migrant sample sizes from both flyways may improve the precision in assigning birds to geographic origin. In addition, obtaining similar results from multiple isotopes over multiple years would provide greater confidence in origin maps for migratory species and are worth pursuing in future studies.

In summary, birds caught along the Pacific Coast Flyway have origins that overlap with those birds caught along the Intermountain Flyway, consistent with the absence of population genetic structure in mitochondrial sequence data among juveniles sampled on these flyways. However, overlapping regions of origin for migrating juveniles does not preclude the possibility of fine-scale population structure among regions, as has been seen in other raptors in western North America [54, 87]. Overall birds that originate from the Rocky Mountain Range of North America appear to choose to travel through either migration route, but discrepancies in the predicted origins based on hydrogen and oxygen isotopes encourage caution, and further studies in how *A. striatus* migrate in western North America.

### Feather isotope composition and life history factors

Our understanding is still poor about why the strong correlation between δ^18^O and δ^2^H values in meteoric (source) water [44] breaks down in the tissues of different groups of vertebrates. Isotopic variability has been observed in δ^2^H_F_ values for wild American Kestrels (*Falco sparverius*) at a local scale [60], and in δ^2^H_F_ and δ^18^O_F_ values in laboratory-controlled groups of House Sparrows (*Passer domesticus*) [77] and Japanese Quail (*Cortunix japonica*) [88]. This variability is thought to be due to differences in diet and water requirements, seasonal timing of water use, metabolism, and evaporative cooling effects, among other factors [88, 89].

A variety of factors may have contributed to the different predictive ability of δ^18^O_F_ and δ^2^H_F_ values in *A. striatus*. Carnivorous animals, including birds, are known to show more positive δ^2^H bone collagen values compared to δ^2^H source water values [90], and hydrogen isotopic composition in organic tissue seems to be influenced more by the diet consumed than the water used, compared to the oxygen isotopic composition in the same tissue [78, 88, 89, 91]. Oxygen isotopic composition may also be affected by atmospheric and dissolved oxygen in body water, in addition to diet and environmental water [89]. In addition, consuming prey from different trophic levels (herbivores vs. insectivores) might affect the hydrogen and oxygen isotopic compositions differently for a raptor such as *A. striatus*. Elevation may also affect the δ^2^H_P_ values [92], and changes in elevation performed during seasonal movement may result in different isotopic compositions then expected [93].Climatic factors may also play a role in the development of different isotopic compositions for raptorial birds from different habitats on the continent. Climate and aridity can have profound effects on the degree of variation (fractionation) of water δ^18^O values measured in animal body water [86, 94, 95], as well as the proportion of drinking versus metabolically produced water in the body-water pool [88]. Moreover, previous work analyzing isotopic information for a variety of raptor species across North America found δ^2^H_F_ enriched values for birds from the northern ranges of the Rocky Mountains in ID and MT compared to δ^2^H_P_ values at the same locations [51]. More work is needed examining the effects of diet and climate on isotopic compositions for birds from different trophic levels to determine if prey, habitat, or both, may play a role in variation in δ^2^H and δ^18^O values.

The age of the animal has also been found to have a significant effect on the δ^2^H_F_ value for birds, with adults showing more positive isotopic compositions relative to younger animals [61, 96–97]. Previous studies on breeding populations of known origin suggest a link between high variability in adult δ^2^H_F_ values, and breeding behavior and physiological effort during the breeding season [60, 62]. Little work has been done using oxygen variation to determine origin of birds from different age classes. While we did not assess the origin of adult *A. striatus* migratory samples, our results for museum samples of known origin are similar between δ^18^O_F_ and predicted δ^18^O_P_ for adult and juvenile *A. striatus* (S3 Fig). Perhaps δ^18^O_F_ values of adult predatory birds may not be affected by the enrichment seen in tissue δ^2^H, possibly because δ^18^O values are more strongly influenced by environmental water then diet [89], and are not as affected by evaporative, metabolic, and respiratory water losses [62].

Research focused on the relationship between the known origin of a bird, its physiological condition, aspects of its behavior, and its diet and drinking regime will no doubt improve our understanding of what “sets” the oxygen and hydrogen isotope composition in feathers. In addition, previous work has identified that the determination of origin for species of various trophic levels is improved when species specific calibration curves for hydrogen are included [48, 98]. Ideally, species specific fractionation factors for hydrogen and oxygen isotopes should be considered along with isotopic variation observed in birds due to differences in age and trophic level. Further research that examines how these variables affect the fractionation factors for hydrogen and oxygen isotope analyses will be essential before we can use hydrogen or oxygen as a reliable tool for examining the origin of all birds. The results from this study on *A. striatus* are encouraging.

### Uncertainties in isoscape modeling methodology

IsoMAP allows users to create and test large scale models of spatial isotopic variation for a specific area of interest. IsoMAP also allows creation of probability density maps to show uncertainty due to sample variability. It is known that predictions based on isoscape modeling methodology might be affected by several factors. The isoscapes created for this study in IsoMAP.com are based on precipitation isotope ratio values collected through a global network of stations [68, 70, 71]. Although the global sampling of precipitation isotope ratios used for the isoscape predictions is spatially and temporally uneven, the coverage in North America is more thorough then in other areas [66]. Therefore, the trends of δ^2^H_P_ and δ^18^O_P_ values can be relatively robust on large scales, but they may not capture the variability present at more limited spatial or temporal scales [67]. This can make attribution challenging or less precise.

Other sources of uncertainty in precipitation isotope values modeled using isoscapes are due to large grid sizes, the integration of isotope data over multiple years, and the interpolation model error. Collection location uncertainty, especially for museum specimens that may have a location description that is not geographically specific, should also be taken into account. Spatial and temporal resolution could be improved by the addition of quality precipitation isotope data to the existing network through platforms like IsoMAP [67].

### Conclusion

Our assignment evaluation demonstrates that the hypothesis that juvenile migratory *A. striatus* birds caught along two distinct migration routes on opposite sides of the Sierra Nevada Mountains of North America (Pacific Coast and Intermountain Migratory Flyways) come from different natal populations can be rejected (Fig 6). We found an overlap in the assigned natal territories of the migrating birds from the two migration routes. Birds captured along the Pacific Coast Flyway had a range of δ^2^H_F_ and δ^18^O_F_ values that were consistent with precipitation found not only along the west coast, but also in the western interior regions in the US and Canada. The birds migrating along the Intermountain Flyway had a more limited geographical origin that also differed if predicted based on SIA of hydrogen or oxygen. The methodology based on oxygen appeared to have less predictive power than the one based on hydrogen perhaps because of the weaker relationships linking feather oxygen isotope ratios to those in precipitation.

We conclude that juvenile migrating *A. striatus* in western North America do not differentiate into fully separate migratory populations. For this difficult-to-track and secretive breeding raptor, our data can provide clues to the origins of birds caught along these two migration routes, and that consideration must be given to both flyways when examining changes in population size at breeding origins, especially along the Rocky Mountain Range and in the interior western regions of the species’ range. However, further work will need to be done, with larger sample sizes, to determine what may be driving the lack of correlation found between the feather hydrogen and oxygen stable isotope compositions of *A. striatus* that migrate through the Intermountain Flyway.

Results from this study corroborate previous work showing that feather isotopes can be useful for identifying sites of origin for migrating birds, but also highlight that caution must be taken when interpreting the outcome, especially if derived by the stable oxygen isotope composition of feathers. Detailed studies on the sources of isotope variation at stages along the path of ingestion and assimilation of water into body tissues, including different trophic levels, life history stages, and geographic complexity, could provide insight for a wider application of SIA of hydrogen and oxygen to track movement of different organisms, and especially wild populations. Identification of natal and breeding habitats has important conservation implications, specifically because the movement of migratory species can often span across large geographic areas and international borders. Organization of conservation efforts for such species requires a precise understanding of movement patterns and connections between breeding sites, migratory pathways, and wintering grounds. The continual development and testing of intrinsic methods, such as stable isotopes, to track animals using feathers or hair can greatly improve our insight into how animals, such as raptors like *A. striatus*, travel and move across their ranges.

## Acknowledgements

This research is an outgrowth of a project for the class *Stable Isotope Ecology* taught at the University of California, Berkeley (by T.E. Dawson and S. Mambelli). We thank Hiromi Uno and Tami Mau for helpful discussion that improved this research, and Paul D. Brooks and Wenbo Yang for their assistance with IRMS analyses. We thank the following museums and migratory sites for all of their hard work collecting and providing us with feathers for this study: CAS, CRCM, MVZ, SDNHM, UWYMV, GGRO, and HawkWatch International Goshutes banding station. This is contribution #141 for the Golden Gate Raptor Observatory.

## Supporting Information

**Table S1: Oxygen stable isotope composition of feathers of adult museum Sharp-shinned Hawk (*Accipiter striatus*) specimens (δ^18^O_f_ values (‰)) (*n* = 25), and isoscape modeled stable hydrogen and oxygen isotope composition of precipitation (δ^18^O_p_ values (‰)) of known breeding origin.**

**Table S2: Stable isotopic compositions of museum feathers and estimates of precipitation isotopes used to test assignment of origin models.** Stable hydrogen (δ^2^H_F_ ‰) and oxygen (δ^18^O_F_ ‰) isotopic composition of feathers for juvenile museum Sharp-shinned Hawk (*Accipiter striatus*) specimens of known natal origin and estimated isotopic compositions of precipitation (δ^2^H_P_ and δ^18^O_P_ ‰) from the transfer functions used to determine assignment of origin. The states where samples were collected and the museum Specimen IDs are included.

**S1 Fig: Map of sample locations for museum specimens of juvenile and adult Sharp-shinned Hawk (*Accipiter striatus*) feathers.** Sampling locations are shown in reference to the species known range in Western North America (light gray), and suitable breeding forest habitat (dark gray). Juveniles samples (*n* = 23) are shown as triangles, and adult samples (*n* = 25) are shown as circles. State and country boundaries are modified from public domain GIS files, US Census Bureau (2016) and Natural Earth (2020). Species range acquired from Birdlife International and NatureServe (2015), and data to create the GIS biome layer acquired from Brown, Bennan, and Unmack (2007).

**S2 Fig: Stable hydrogen (δ^2^H_P_ ‰) and oxygen (δ^2^O_P_ ‰) isoscapes created within IsoMAP.** These were used to determine transfer functions for specimens of known origin and performing assignment of origin for migrating specimens. The hydrogen isoscape (A) produced was based on 117 stations, had resolution of 9×9 km, a correlation parameter of 0.93, and included the variables elevation (ETOPO, *P* < 0.001), latitude (*P* < 0.001) and longitude (*P* = 0.06) (available as IsoMAP job key 50333 (Marrack 2015)). The most robust oxygen isoscape (B) was based on 120 stations, had resolution of 9×9 km, a correlation parameter of 0.92, and included the variables elevation (ETOPO, *P* < 0.001), latitude (*P* < 0.001) and longitude (*P* = 0.05) (available as IsoMAP job key 63026 (Marrack 2017)).

**S3 Fig: Relationship between the stable isotopic compositions of museum feathers and the isoscape model of precipitation for adult and juvenile Sharp-shinned Hawks (*Accipiter striatus*).** Stable oxygen (δ^18^O_F_ ‰) isotopic composition of feathers for museum Sharp-shinned Hawk (*Accipiter striatus*) specimens of known natal/breeding origin and isoscape modeled isotopic composition of precipitation (δ^18^O_P_ ‰) at the collection locations of (a) juvenile (black dots) birds as well as (b) adult birds (white dots) (*n* = 48). The linear regression for both juvenile and adult birds is *y* = 0.29 *x* – 14.77, R^2^ = 0.1, *P* = 0.03 (dashed line). Note that this equation is similar in slope and intercept to the linear regression relationship for juvenile birds (*y* = 0.385*x* – 14.67 (R^2^ = 0.14, *P* = 0.07) (solid line)).

**S4 Fig: Relationship between the stable isotopic compositions of museum feathers and the isoscape model of precipitation.** Stable hydrogen (δ^2^H_F_ ‰) and oxygen (δ^18^O_F_ ‰) isotopic composition of feathers for museum juvenile Sharp-shinned Hawk (*Accipiter striatus*) specimens (*n* = 23) of known natal origin and the isoscape modeled isotopic compositions of precipitation (δ^2^H_F_ and δ^18^O_P_ ‰) at the collection locations: (a) δ^2^H_F_ values of birds versus δ^2^H_P_ values, (b) δ^18^O_F_ values of birds versus δ^18^O_P_ values.

**S5 Fig: Probability density maps predicting the origin of museum specimens of Sharp-shinned Hawks (*Accipiter striatus*) with known collection locations.** Maps are based on predicted δ^2^H_P_ values (‰) (left panels) and predicted δ^18^O_P_ values (‰) (right panels) for birds captured at known locations. Each map represents the probability density surface created for an individual bird with the known sampling location shown by a circle. State and country boundaries are from public domain GIS files US Census Bureau (2016) and Natural Earth (2020). Species range acquired from BirdLife International and NatureServe (2015), and data to create the GIS biome layer acquired with permission from Brown, Bennan, and Unmack (2007).

