## Supplementary figures and images for "Using oxygen and hydrogen stable isotopes to track the migratory movement of Sharp-shinned Hawks (*Accipiter striatus*) along Western Flyways of North America"

### S1 Fig

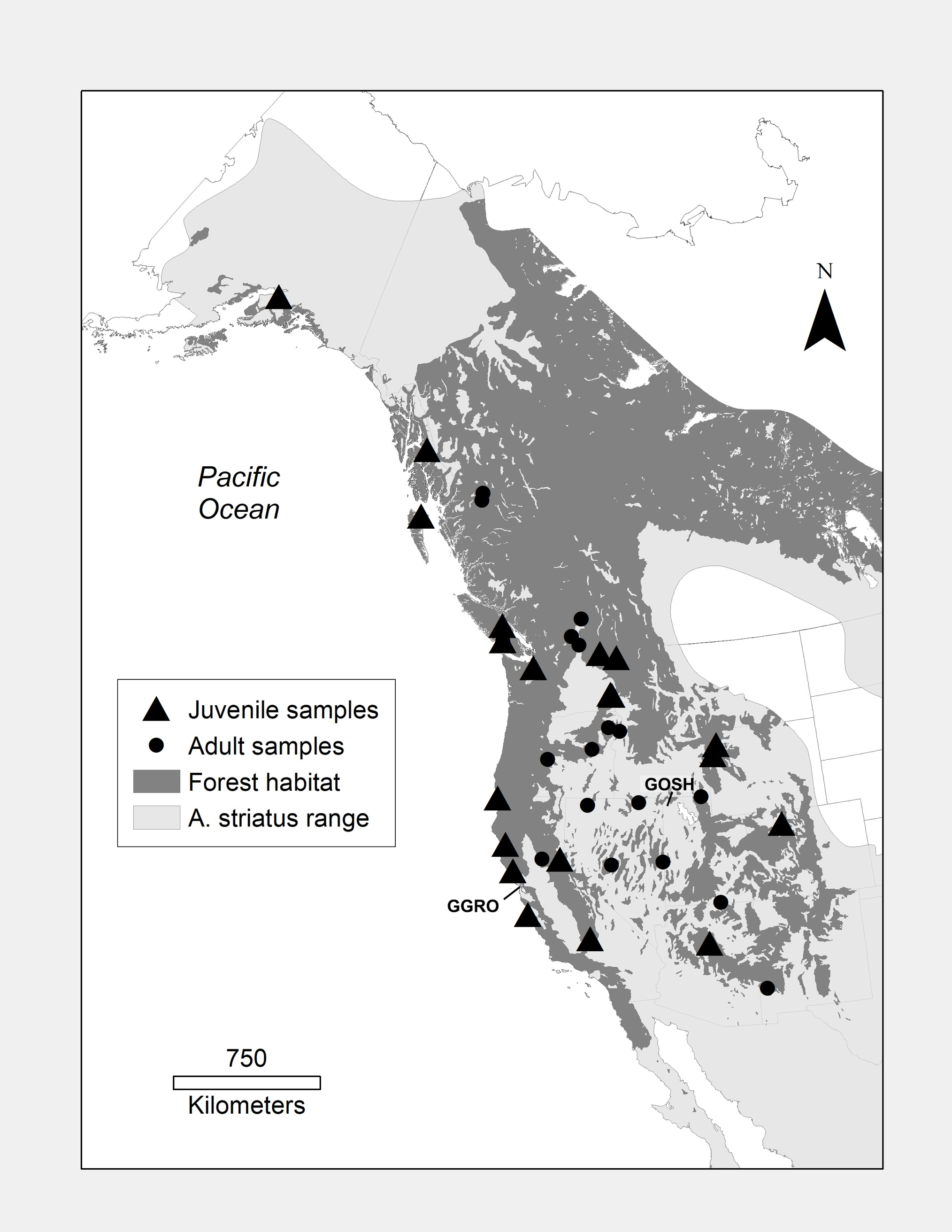

### S3 Fig

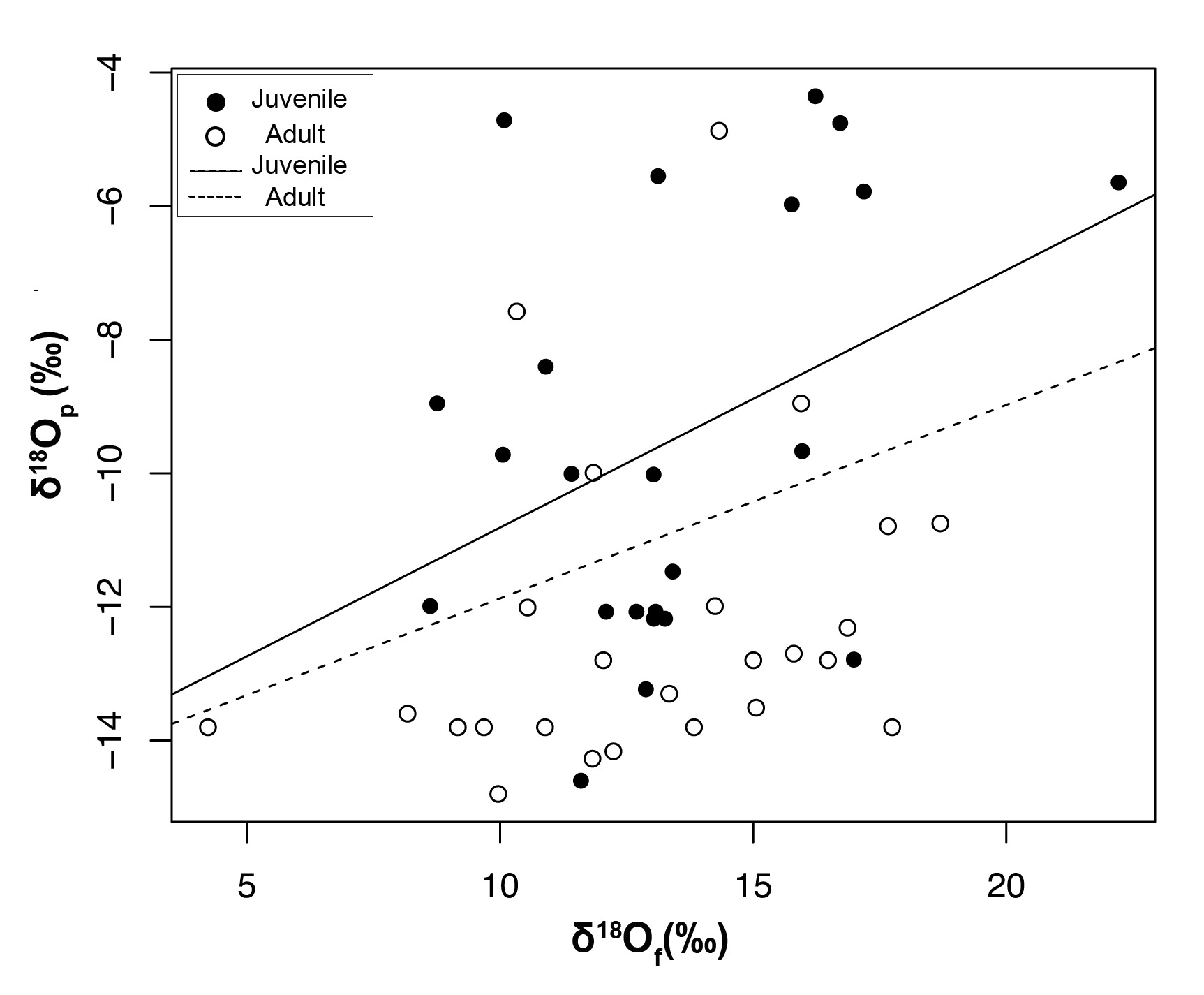

### S4 Fig

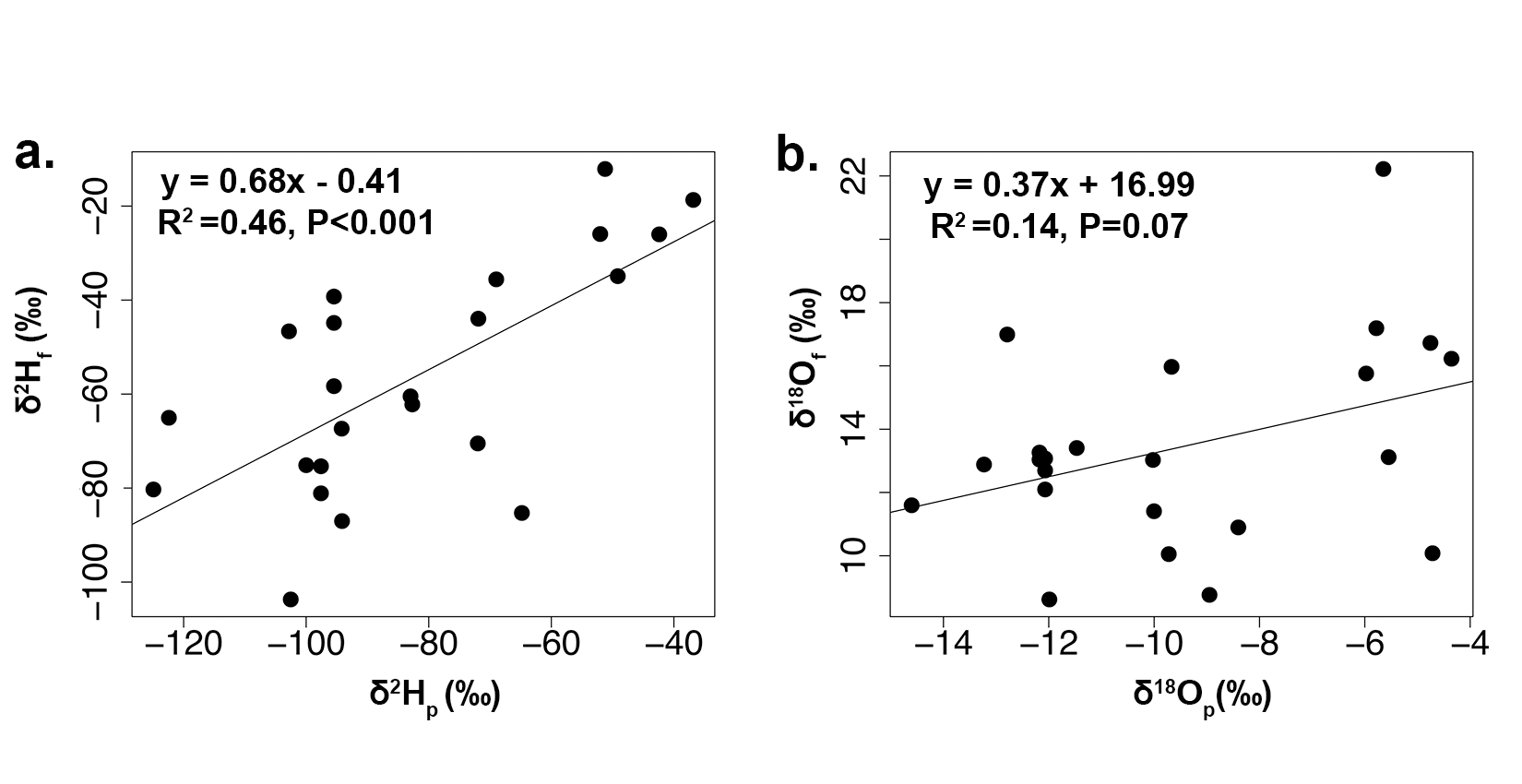
