## Supplementary material for "Using oxygen and hydrogen stable isotopes to track the migratory movement of Sharp-shinned Hawks (*Accipiter striatus*) along Western Flyways of North America": S2 Fig

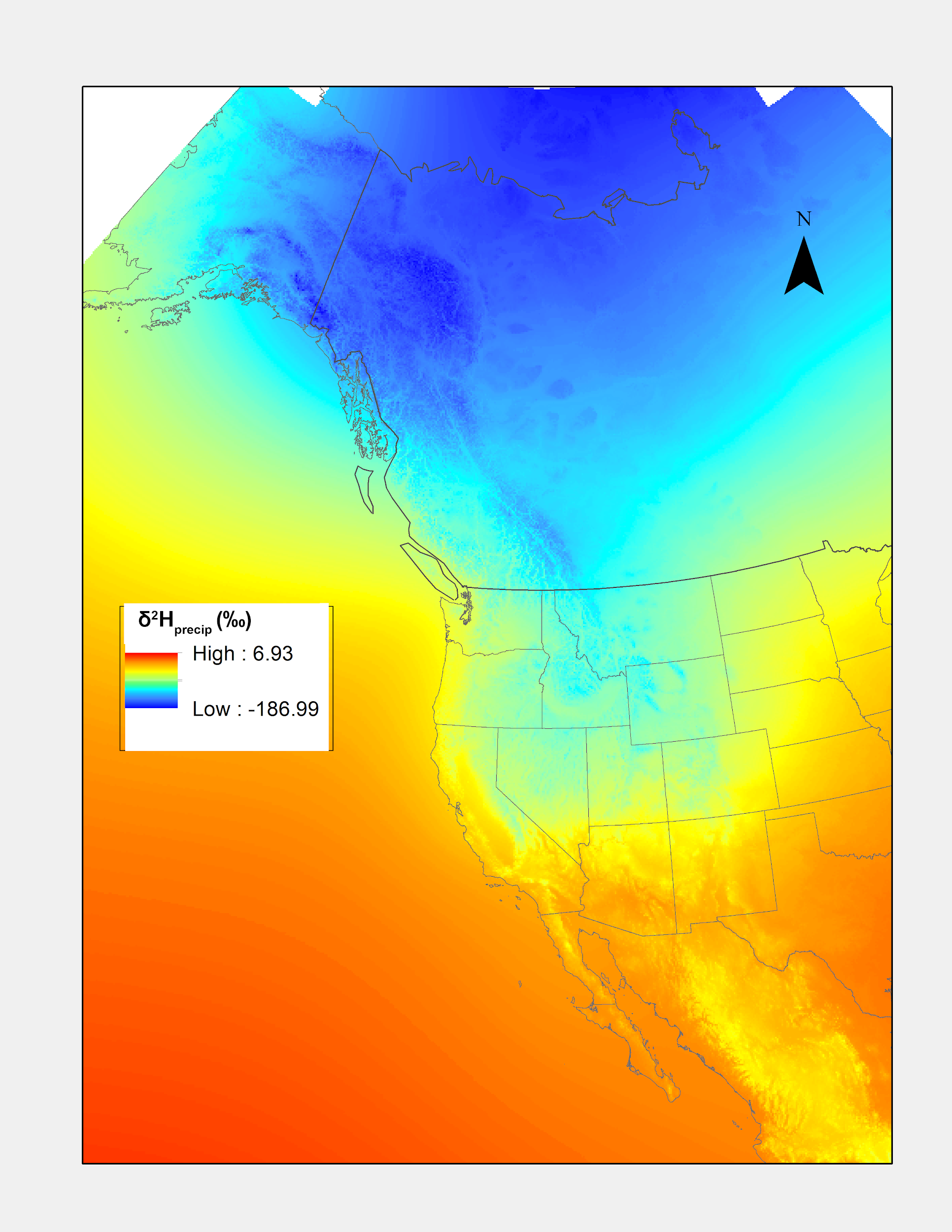

**A**

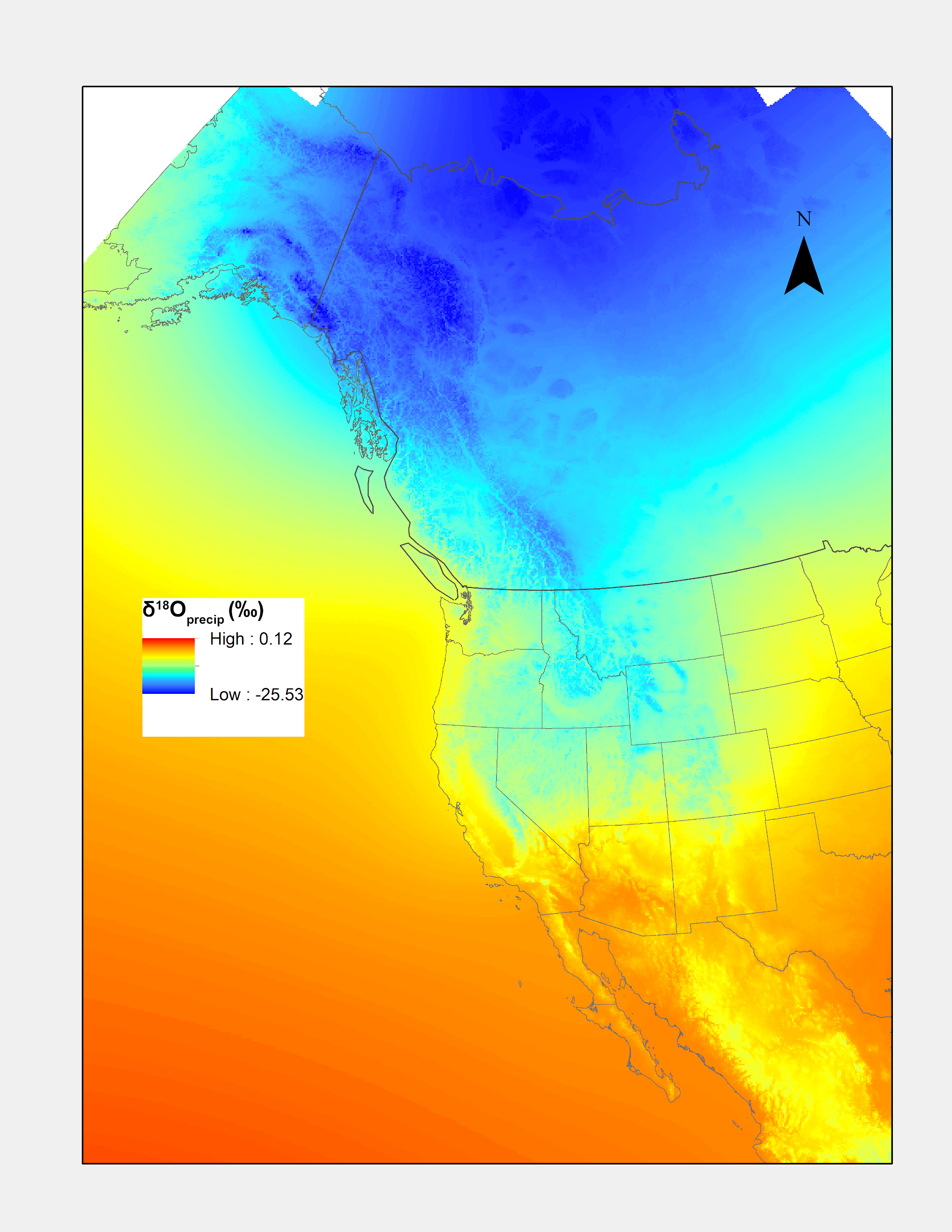

**B**

**S2 Fig: Stable hydrogen (δ^2^H_P_ ‰) and oxygen (δ^18^O_P_ ‰) isoscapes created within IsoMap.** These were used to determine transfer functions for specimens of known origin and performing assignment of origin for migrating specimens. The hydrogen isoscape (A) produced was based on 117 stations, had resolution of 9x9 km, a correlation parameter of 0.93, and included the variables elevation (ETOPO, P < 0.001), latitude (P < 0.001) and longitude (P = 0.06) (available as IsoMAP job key 50333 (Marrack 2015)). The most robust oxygen isoscape (B) was based on 120 stations, had resolution of 9x9 km, a correlation parameter of 0.92, and included the variables elevation (ETOPO, P < 0.001), latitude (P < 0.001) and longitude (P = 0.05) (available as IsoMAP job key 63026 (Marrack 2017)).
