## Supplementary material for "Using oxygen and hydrogen stable isotopes to track the migratory movement of Sharp-shinned Hawks (*Accipiter striatus*) along Western Flyways of North America": S5 Fig

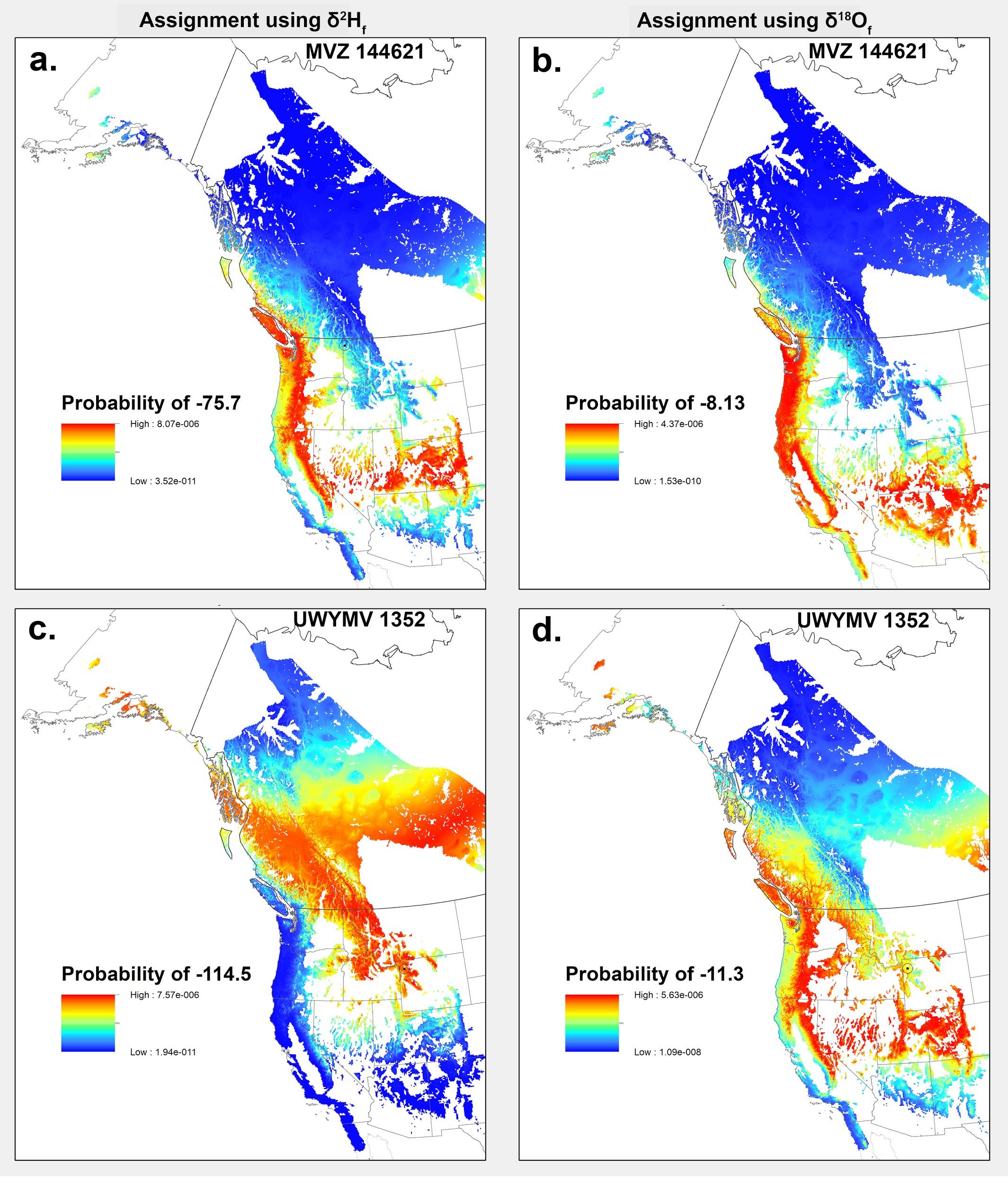

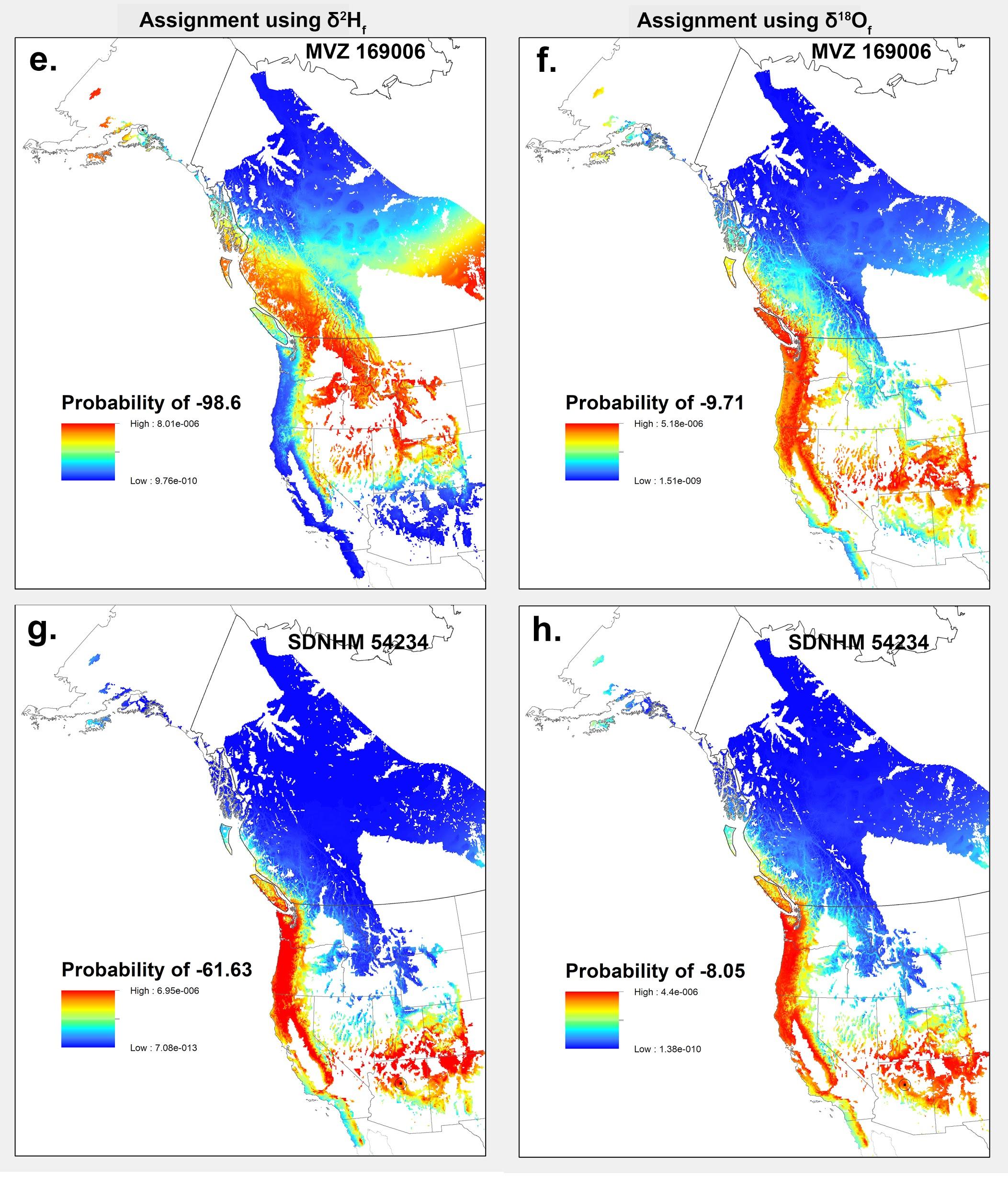

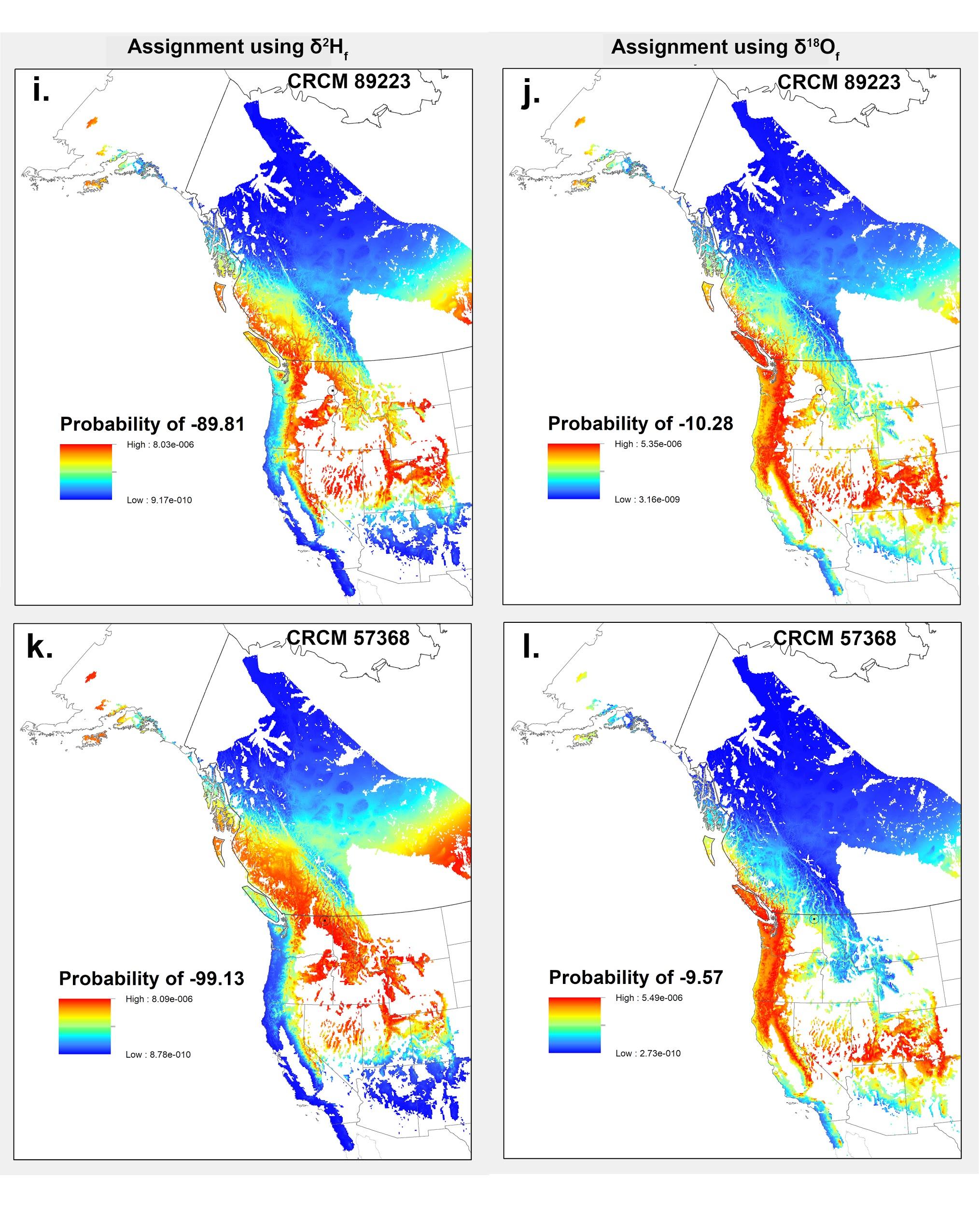

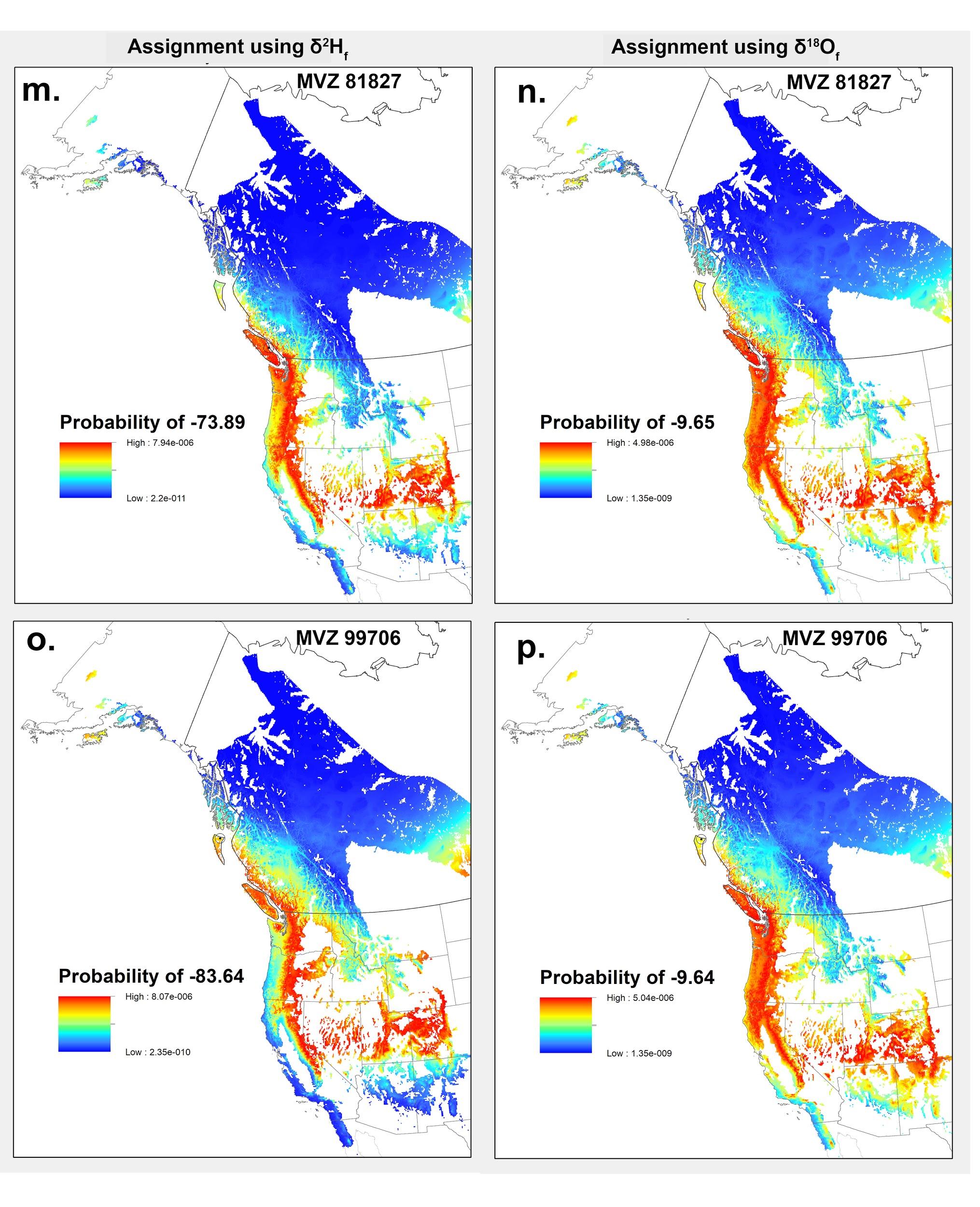

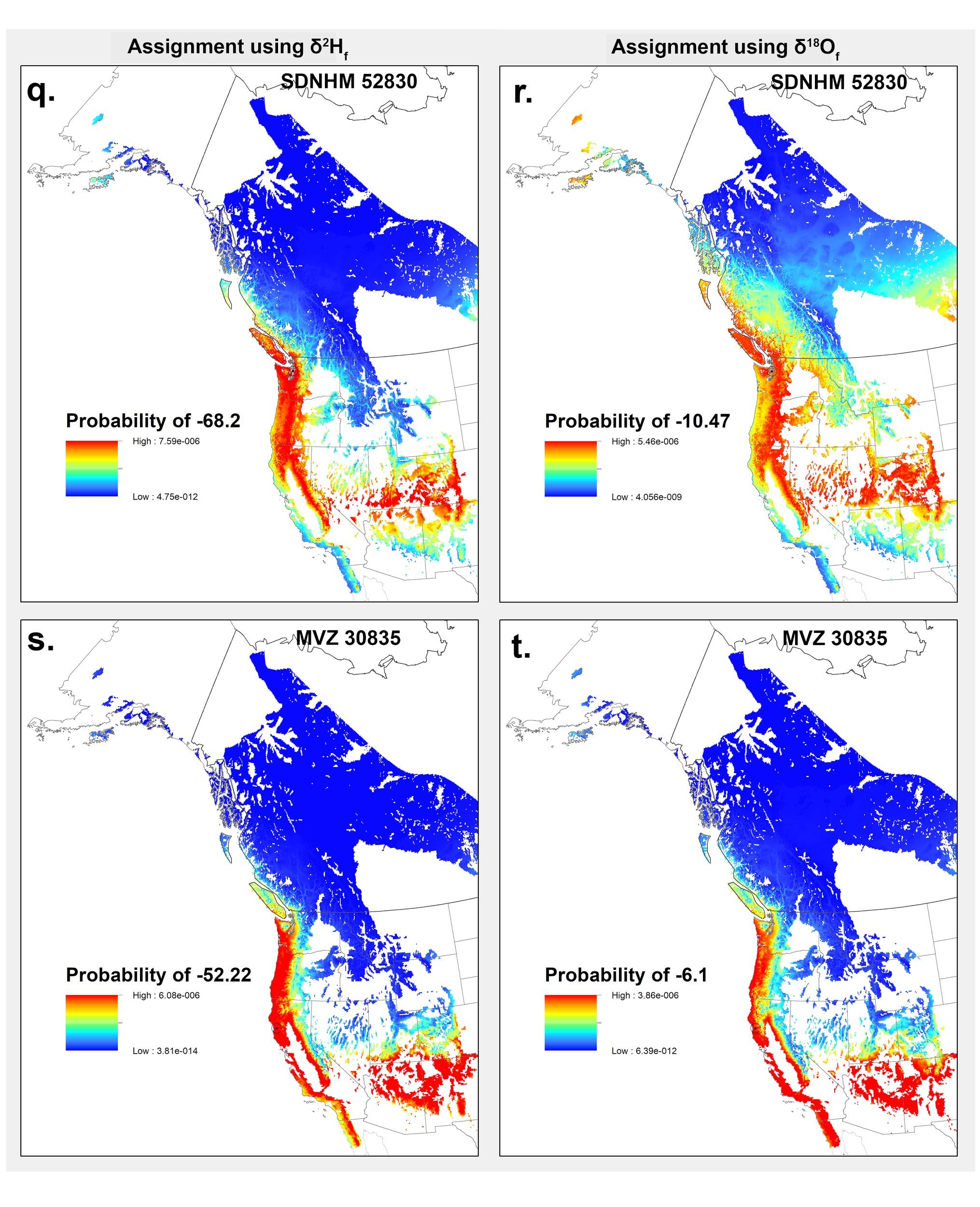

**S5 Fig. Probability density maps predicting the origin of museum specimens of Sharp-shinned Hawks (*Accipiter striatus*) with known collection locations.** Maps are based on predicted δ^2^H_P_ values (‰) (left panels) and predicted δ^18^O_P_ values (‰) (right panels) for birds captured at known locations. Each map represents the probability density surface created for an individual bird with the known sampling location shown by a circle. State and country boundaries are from public domain GIS files US Census Bureau (2016) and Natural Earth (2020). Species range acquired from BirdLife International and NatureServe (2015), and data to create the GIS biome layer acquired with permission from Brown, Bennan, and Unmack (2007).
