## Supplementary material for "Using oxygen and hydrogen stable isotopes to track the migratory movement of Sharp-shinned Hawks (*Accipiter striatus*) along Western Flyways of North America": Table S1

**STable 1:** Oxygen stable isotope composition of feathers of adult museum Sharp-shinned Hawk (*Accipiter striatus*) specimens (δ^18^O_f_ values (‰)) (*n* = 25), and isoscape modeled stable hydrogen and oxygen isotope composition of precipitation (δ^18^O_p_ values (‰)) of known breeding origin.

| **^a^Museum** | **^b^Specimen No.** | **Life stage** | **^c^State** | **Latitude** | **Longitude** | **δ^18^O_f_ (‰)** | **δ^18^O_p_ (‰)** |
| --- | --- | --- | --- | --- | --- | --- | --- |
| MVZ | 9094 | Adult | NV | 41.67278 | -118.6275 | 16.48 | -12.8 |
| MVZ | 9095 | Adult | NV | 41.67278 | -118.6275 | 15.8 | -12.7 |
| MVZ | 15621 | Adult | BC | 48.9895 | -124.8 | 11.84 | -9.99 |
| MVZ | 42040 | Adult | BC | 55.25 | -127.6667 | 11.82 | -14.27 |
| MVZ | 42042 | Adult | BC | 55.5825 | -127.6667 | 9.96 | -14.8 |
| MVZ | 53345 | Adult | CA | 39.1144136 | -121.26609 | 10.33 | -7.58 |
| MVZ | 57317 | Adult | NV | 38.92 | -117.16222 | 17.67 | -10.79 |
| MVZ | 58092 | Adult | UT | 41.94694 | -111.62568 | 13.34 | -13.3 |
| MVZ | 64641 | Adult | NV | 39.01722 | -114.10722 | 18.7 | -10.75 |
| MVZ | 71811 | Adult | OR | 43.7147 | -121.2136 | 14.25 | -11.99 |
| MVZ | 73402 | Adult | OR | 44.2483 | -118.4086 | 8.17 | -13.6 |
| MVZ | 73403 | Adult | OR | 45.2558 | -117.3669 | 12.24 | -14.16 |
| MVZ | 81825 | Adult | BC | 49.0333 | -119.45 | 10.54 | -12.01 |
| MVZ | 99714 | Adult | BC | 49.4167 | -120 | 13.83 | -13.8 |
| MVZ | 99715 | Adult | BC | 49.4167 | -120 | 4.23 | -13.8 |
| MVZ | 99716 | Adult | BC | 49.4167 | -120 | 9.16 | -13.8 |
| MVZ | 99717 | Adult | BC | 49.4167 | -120 | 10.89 | -13.8 |
| MVZ | 99718 | Adult | BC | 49.4167 | -120 | 9.68 | -13.8 |
| MVZ | 99723 | Adult | BC | 49.4167 | -120 | 17.75 | -13.8 |
| MVZ | 99727 | Adult | BC | 50.2333 | -119.35 | 15 | -12.8 |
| MVZ | 99728 | Adult | BC | 50.2333 | -119.35 | 12.04 | -12.8 |
| MVZ | 99740 | Adult | NM | 32.9634798 | -108.61215 | 14.33 | -4.87 |
| MVZ | 144623 | Adult | ID | 45.0884 | -116.6367 | 16.86 | -12.31 |
| MVZ | 144628 | Adult | NV | 41.795 | -115.48778 | 15.06 | -13.51 |
| MVZ | 64967 | Adult | UT | 37.03475 | -110.86898 | 15.95 | -8.95 |

^a^Museum: MVZ = Museum of Vertebrate Zoology, University of California, Berkeley, CA, USA

^b^Specimen: Full information on each specimen can be obtained by taking the specimen number and searching for it in the online databases Vertnet.org and <https://arctosdb.org/>

^c^State or Provinces: BC = British Colombia, Canada; CA = California, USA; ID = Idaho, USA; NM = New Mexico, USA; NV = Nevada, USA; OR = Oregon, USA; UT = Utah, USA
