## Supplementary material for "Using oxygen and hydrogen stable isotopes to track the migratory movement of Sharp-shinned Hawks (*Accipiter striatus*) along Western Flyways of North America": Table S2

**Table S2: Stable isotopic compositions of museum feathers and estimates of precipitation isotopes used to test assignment of origin models.** Stable hydrogen (δ^2^H_F_ ‰) and oxygen (δ^18^O_F_ ‰) isotopic composition of feathers for juvenile museum Sharp-shinned Hawk (*Accipiter striatus*) specimens of known natal origin and estimated isotopic compositions of precipitation (δ^2^H_P_ and δ^18^O_P_ ‰) from the transfer functions used to determine assignment of origin. The states where samples were collected, and the museum Specimen IDs are included.

| **Hydrogen Transfer Function** | **^a^Museum Sample ID** | **Measured δ^2^H_F_** | **Estimated δ^2^H_p_** | **^b^Locality** |
| --- | --- | --- | --- | --- |
| δ^2^H_p_=(δ^2^H_F_*0.68)-43.98, RSE 18.3 | MVZ 99706 | -58.3 | -83.64 | BC |
|  | MVZ 169006 | -80.3 | -98.6 | AK |
|  | UWYMV 1352 | -103.7 | -114.5 | WY |
|  | CRCM 57-368 | -81.1 | -99.13 | WA |
|  | MVZ 30835 | -12.1 | -52.22 | CA |
|  | SDNHM 52830 | -35.6 | -68.20 | WA |
|  | MVZ 144621 | -46.7 | -75.7 | ID |
|  | MVZ 81827 | -44.0 | -73.89 | BC |
|  | SDNHM 54234 | -26.0 | -61.63 | AZ |
|  | CRCM 89-223 | -67.4 | -89.81 | WA |

| **Oxygen Transfer Function** | **^a^Museum Sample ID** | **Measured δ^18^O_F_** | **Estimated δ^18^O_p_** | **^b^Locality** |
| --- | --- | --- | --- | --- |
| δ^18^O_p_=(δ^18^O_F_*0.385)-14.67, RSE 3.0 | MVZ 30835 | 22.22 | -6.12 | CA |
|  | UWYMV 1352 | 8.62 | -11.35 | WY |
|  | CRCM 57-368 | 13.26 | -9.57 | WA |
|  | MVZ 99706 | 13.07 | -9.64 | BC |
|  | MVZ 169006 | 12.88 | -9.71 | AK |
|  | SDNHM 52830 | 10.90 | -10.47 | WA |
|  | MVZ 144621 | 16.99 | -8.13 | ID |
|  | MVZ 81827 | 13.03 | -9.65 | BC |
|  | SDNHM 54234 | 17.19 | -8.05 | AZ |
|  | CRCM 89-223 | 11.41 | -10.28 | WA |

^a^Museums: MVZ = CRCM = Charles R. Connor Museum; Museum of Vertebrate Zoology; SDNHM = San Diego Natural History Museum; UWYMV = University of Wyoming Museum of Vertebrates

^b^States and Provinces: AK = Alaska, USA; AZ = Arizona, USA; BC = British Columbia, Canada; CA = California, USA; ID = Idaho, USA; WA = Washington, USA; WY = Wyoming, USA
